# Locally acting transcription factors are required for p53-dependent *cis-*regulatory element activity

**DOI:** 10.1101/761387

**Authors:** Allison N. Catizone, Gizem Karsli Uzunbas, Petra Celadova, Sylvia Kuang, Daniel Bose, Morgan A. Sammons

## Abstract

The master tumor suppressor p53 controls transcription of a wide-ranging gene network involved in apoptosis, cell cycle arrest, DNA damage repair, and senescence. Recent studies revealed pervasive binding of p53 to *cis-*regulatory elements (CRE), which are non-coding segments of DNA that spatially and temporally control transcription through the combinatorial binding of local transcription factors (TFs). Although the role of p53 as a strong *trans-*activator of gene expression is well known, the co-regulatory factors and local sequences acting at p53-bound CREs are comparatively understudied. We designed and executed a massively parallel reporter assay (MPRA) to investigate the effect of transcription factor binding motifs and local sequence context on p53-bound CRE activity. Our data indicate that p53-bound CREs are both positively and negatively affected by alterations in local sequence context and changes to co-regulatory TF motifs. We identified a SP1/KLF family motif located in an intronic p53 CRE that is required for the endogenous expression of the p53-dependent gene *CCNG1.* We also identified ATF3 as a factor that co-regulates the expression of the p53-dependent gene *GDF15* through binding with p53 in an upstream CRE. Loss of either p53 or ATF3 severely reduces CRE activity and alters endogenous *GDF15* mRNA levels in the cell. Our data suggests that p53 has the flexibility to cooperate with a variety of transcription factors in order to regulate CRE activity. By utilizing different sets of co-factors across CREs, we hypothesize that p53 activity is guarded against loss of any one regulatory partner allowing for dynamic and redundant control of p53-mediated transcription.

## Introduction

The master tumor suppressor p53 is a transcription factor with key roles in preserving genome fidelity and cellular homeostasis. In support of these activities, p53 regulates a core transcriptional program involved in cellular processes like cell cycle arrest, apoptosis, DNA repair, and senescence (1–3). Loss of p53 activity is strongly linked to increased cancer risk and decreased life expectancy, and misregulation of p53 is associated with numerous other human disorders. Recent estimates suggest that p53 is mutated in greater than 30% of cancer cases and the majority of p53 variants are unable to bind DNA and to enact a tumor suppressive gene expression program (4). The mechanisms by which tumorigenesis progresses in the presence of wild type p53 activity have not been well characterized. Recent evidence suggests that sequence variation within *cis-*regulatory elements (CREs) can influence p53 binding, transcriptional activity, and tumor suppressor function (5–7). The critical nature of the core p53 response element (p53RE) on p53 binding and CRE activity is well understood (8–10), but the influence of local sequence variation and the role of transcription factor motifs outside of the p53RE on p53 activity remains an open and vital question.

*Cis-*regulatory elements, such as promoters and enhancers, govern gene expression through temporal, spatial, and quantitative control of transcription (11, 12). While multiple models for CRE function have been proposed, the majority involve cooperative binding and activity of multiple transcription factors and cofactors acting locally to fine-tune gene expression (11–13). The presence and availability of transcription factors, repressors, and other cofactors vary across cell states such as development, stress, disease, and cell type (14–17). This variability provides a mechanism for differential CRE activity and downstream gene expression. Loss of transcription factor binding within a CRE, through variation in DNA sequence or through changes in *trans-*factor availability, can strongly influence CRE activity and gene expression (11, 12, 18), with direct implications in numerous developmental and disease states (14, 16).

While general transcription factors, like the TFIID complex (19, 20), are involved in p53-dependent trans-activation at promoters, the requirement for other sequence-specific trans-factors at distally-acting CREs is unknown. A novel model was recently proposed whereby binding of a single transcription factor, in this case p53, was necessary and sufficient for CRE activity (9). This model was supported by another study suggesting that a canonical p53 response element (p53RE) is the only sequence-based determinant of p53-dependent CRE activity (10). However, multiple p53-dependent CREs have been reported to require other locally-acting transcription factors, in line with established CRE mechanisms like the enhanceosome and billboard models (18, 21). For example, CRISPR/Cas9-based screening identified a CEBPβ-binding site within a CRE regulating *CDKN1A*/p21 that required for p53-dependent senescence (22). Transcription factors such as those in the AP-1 family and SP1 have also been implicated in the activation of p53-dependent gene targets (23–25).

In order to directly address whether additional cofactors are required for p53-dependent transcriptional activity, we examined the effect of local sequence variation on putative p53 CREs using a massively parallel reporter assay (MPRA). Our results suggest that sequences flanking p53RE and transcription factors other than p53 act within CREs to facilitate p53-dependent transcriptional activation. Consistent with previous reports, the p53RE is a strong determinant of p53-induced activity. Loss of p53 occupancy through sequence manipulation or depletion of p53 protein strongly reduces CRE activity. We also identified sequences outside of the p53RE that positively or negatively regulate transcription. This includes a conserved SP1/KLF family binding site required for optimal transcription of the p53-dependent gene *CCNG1 (*cyclin G1). We also identified two distinct CREs required for p53-dependent transcriptional activation of *GDF15*, a gene recently identified as a key mediator of inflammation and metabolic function. Importantly, each CRE requires a unique cohort of transcription factors for optimal activity, suggesting p53-bound CREs use different combinations of transcription factors to drive p53-dependent transcription. Thus, indicating p53’s flexibility to utilize various, locally available transcription factors to drive expression of a key tumor suppressor network.

## Materials and Methods

### Cell Culture

HCT116 parental, TP53-/-, and ATF3-/- lines were cultured in McCoy’s 5A media with 10% fetal bovine serum. HCT116 ATF3-/- cell line was a kind gift of Chunhong Yan (Augusta University) (26). MCF10A cells were grown in HuMEC media (Gibco). Mouse Embryonic Fibroblasts (MEFs) were grown in DMEM with 10% FBS and were a kind gift of Jing Huang (National Cancer Institute, NIH). All cell lines were cultured at 37°C and 5% CO_2_ in a water-jacketed incubator.

### Selection of candidate enhancers

Candidate enhancer regions were selected starting with regions of the hg19 genome assembly containing DNAse Hypersensitive Sites (DHS) (wgEncodeRegDnaseClusteredV3 downloaded from http://hgdownload.cse.ucsc.edu/goldenPath/hg19/encodeDCC/wgEncodeRegDnaseClustered/). DHS were then filtered by the presence of a p53 family motif using gimmeMotifs (27). Ultimately, 296 DHS containing a p53 family motif and falling within 100kb of a coding sequence transcriptional start site (TSS) were randomly selected for MPRA analysis. In parallel, 196 enhancers from the FANTOM Ubiquitous Enhancer group were selected as positive controls, with the central 100bp segment of each enhancer used in the MPRA (http://enhancer.binf.ku.dk/presets/Ubiquitous_enhancers_S9.bed). Candidate p53-bound enhancers were shortened to 100bp with the 20bp p53 response element motif at the center with 40bp of flanking genomic context on each side. Each candidate p53-bound enhancer was scrambled in 20bp sections from 5’ to 3’ across the entire length producing. Nucleotide randomization preserved GC content and was performed using EMBOSS shuffleseq (28). As negative controls for regulatory activity, the entire 100bp sequence for all candidate or ubiquitous enhancers was scrambled while preserving GC content.

### Massively Parallel Reporter Assay (MPRA) Oligo Design

Our MPRA was designed using a previously published method (29). In brief, each candidate 100bp regulatory sequence was coupled to 5 separate 12 nucleotide unique molecular identifier (UMI) sequences. Replicates of the test sequences plus controls totaled the library at 12, 035 unique oligos in the orientation of: a 5’ Primer binding overlap, a 100bp enhancer sequence, a spacer for restriction enzyme sites EcoRI and SbfI, a unique enhancer associated 12bp barcode, and a 3’ Primer binding overlap. All sequences for the MPRA oligo library are found in Supplemental Table 1. The final 12, 035 unique oligo pool was synthesized by CustomArray.

### Two-step Vector Library Cloning and Verification

The MPRA lentiviral vector pLs-mP was a gift from Nadav Ahituv (Addgene plasmid # 81225; http://n2t.net/addgene:81225; RRID:Addgene_81225). pLs-mP was digested with the restriction enzymes EcoRI and SbfI yielding two fragments representing the plasmid backbone and the minimal promoter/eGFP. The candidate enhancer pool was PCR amplified using primers SL468 and SL469 in 3 separate reactions of 50ng at 21 cycles each, gel purified, and combined (Supplemental Table 2, cloning_primers tab). The resulting PCR product was ligated to the EcoRI-SbfI digested pLs-mP backbone in 3 separate Gibson assembly reactions (HiFi DNA Assembly, NEB). 2 µL of each Gibson assembly reaction were transformed into Stbl4 electrocompetent *E. coli* (Invitrogen) in 3 separate transformation reactions (1200 V, 200 Ω, BioRad). Transformation reactions were plated on 10 separate 15cm LB-agar plates with 100ug/mL ampicillin selection at 30°C for 48 hours. Colonies were isolated directly from plates and plasmid DNA was individually prepped (ZymoPURE II Midi Plasmid Kit). The resulting DNA was combined to create the Step 1 library. This plasmid pool was then digested with EcoRI and SbfI in three separate reactions at 1 µg each and gel purified. The 780 bp fragment from the original EcoRI-SbfI reaction (containing the minimal promoter/EGFP fragment), was ligated into the digested Step 1 plasmid using T4 ligase. Ligation products were transformed as above and the resulting MPRA library was sequence verified by Illumina sequencing.

### MPRA virus production and transduction

HEK293FT cells were used for virus preparation and were cultured in DMEM with 10% FBS without antibiotic. To make virus, 4 x 10^6^ cells were seeded in 10cm plates 24 hours before transfection. Per 10cm plate, cells were transfected with 8ug of MPRA lentiviral backbone, 4ug of M2G helper plasmid and 8ug of ps-PAX helper plasmid using JetPrime transfection reagent according to manufacturer’s recommendations. Virus was collected from supernatant at 24 and 48 hours, pooled, and filtered using 0.45μm syringe filters. HCT116 TP53^+/+^ and TP53^-/-^ cell lines were seeded 24 hours before viral transduction in three 10cm plates at a concentration of 2.0 x 10^6^ cells/plate. Virus supernatant was combined with 8 µg/mL polybrene, added to the seeded cells, and incubated for 48 hours. Cells were then treated for 6 hours with DMSO (as a control) or 5µM Nutlin-3A to induce p53 activity. One plate was left untreated as the infection control plate for genomic DNA isolation. After 6 hours of treatment, cells were collected in ice-cold 1X PBS, snap-frozen on liquid nitrogen, and stored at −80°C until analysis. Three biological replicates were performed for each condition.

### Amplicon Enrichment and RNA-seq Library Preparation

Total RNA was isolated from DMSO or Nutlin-3A treated cells (EZ RNA Kit, Omega Biotek) with on-column DNAseI treatment. 6ug of resulting total RNA was then taken through an additional round of TurboDNase treatment (ThermoFisher) to ensure complete removal of contaminating genomic DNA. The resulting RNA was split into three 1^st^ strand reverse transcription reactions each using custom barcoded primers to identify the cell line, treatment, and replicate number (Supplemental Table 2, Barcode_primers tab). All 3 cDNA reactions were combined and taken through a 2-step PCR amplification process. In Round 1, each cDNA sample was amplified in 22 separate PCR reactions of 3 cycles each using barcoded primers (Supplemental Table 2). PCR reactions were then pooled and purified using 2.5x AMPure XP beads (Beckman-Coulter). The purified Round 1 PCR product went through a second round of PCR in 8 reactions for 15 cycles each using barcoded primers as described in Supplemental Table 2. Step 2 PCR product was run on a 2% agarose gel and gel purified. Genomic DNA controls were prepared in a similar manner. 500ng of genomic DNA was PCR amplified across 16 separate reactions of 3 cycles each using barcoded primers described in Supplemental Table 2. The pooled PCR product was combined and purified using 2.5X volume AMPure XP beads. The resulting purified DNA was then separated into 16 separate PCR reactions of 15 cycles each, pooled, and gel purified. After the 2-step PCR reaction, DNA amplicons representing the expressed mRNA barcode and the genomic DNA infection control within each biological replicate were combined at equal molarity. An Illumina-compatible sequencing library was generated (NEBNext Ultra II DNA Library Kit, New England Biolabs) for each biological replicate and sequenced using the NextSeq 500 at the University at Albany Center for Functional Genomics.

### Massively Parallel Reporter Assay (MPRA) Data Analysis

Sequencing reads for each individual experimental condition were flanked by a unique 5’ and 3’ amplicon barcode to allow separation from the pooled raw sequencing reads. Barcoded, experimental condition-specific reads were separated into individual files for further analysis using the FastX toolkit (fastx-barcode-splitter, http://hannonlab.cshl.edu). Barcodes for within-pool sequence identification can be found in Supplemental Table 2 (barcode_primers tab). The number of reads containing unique enhancer identifying sequences were then parsed and counted using fastX-collapser from the FastX-toolkit. Raw read counts for each enhancer sequence across cell lines, treatment conditions, and biological replicates be found in Supplemental Table 3. Differential enhancer activity across experimental conditions and cell lines was calculated from raw enhancer barcode read counts using DESeq2 (30). To account for differences in representation across the original viral enhancer sequence library, raw enhancer barcode read counts scaled to transcripts per million (TPM) and were then normalized to the read counts from genomic DNA (fold-change, RNA barcode/ DNA barcode). Normalized enhancer count values for mutant enhancer sequences were then compared to wild-type values using a one-way ANOVA with a post-hoc Tukey HSD test implemented in *R* (31).

### In vivo CRISPR/Cas9 mutagenesis and amplicon ChIP-sequencing

HCT116 colon carcinoma cells were transduced with lentiCas9-Blast and cells stably expressing wild-type spCas9 were selected using 2µg/mL blasticidin. LentiCas9-Blast was a gift from Feng Zhang (Addgene plasmid #52962; http://n2t.net/addgene:52962; RRID:Addgene_52962). *Streptomyces pyogenes* (sp) guide RNA sequences (Table 2) were cloned into the LentiGuide-Puro plasmid (plasmid #52963, Addgene). lentiGuide-Puro was a gift from Feng Zhang (Addgene plasmid # 52963; http://n2t.net/addgene:52963; RRID:Addgene_52963). Both LentiGuid-Puro and LentCas9-Blast were originally published in (32). Viral particles from LentiGuide-Puro were made by transfecting HEK293FT cells with the packaging plasmids psPax2 and pMD2.G and the respective guide RNA viral backbone cloned into pLS-mP. psPAX2 was a gift from Didier Trono (Addgene plasmid # 12260; http://n2t.net/addgene:12260; RRID:Addgene_12260). pMD2.G was a gift from Didier Trono (Addgene plasmid # 12259; http://n2t.net/addgene:12259; RRID:Addgene_12259).

HCT116 cells stably expressing spCas9 were selected with 2 μg/mL puromycin 48 hours after infection. The heterogenous cell pool was then treated for 6 hours with either DMSO or 5µM Nutlin-3A for either qPCR-mediated gene expression measurements or for ChIP. For ChIP, 10 million cells were crosslinked on plate with 1% formaldehyde for 10 minutes at room temperature until the reaction was quenched with 2.5% glycine. Crosslinked cells were processed using standard lysis procedures and chromatin was sonicated using a probe sonicator (Qsonica) at 25% amp for 10 pulses total; 10 seconds on, 50 seconds off for a total of 10 minutes at 4°C. Sheared chromatin was then used in a ChIP assay for p53 (clone DO1, BD Biosciences). 50ng of purified DNA per experimental sample was used in a barcoding PCR reaction (Supplemental Table 2, barcode_primers tab), and amplicons were then used as template for created Illumina-compatible sequencing libraries (NEBNext Ultra II DNA Library Kit). Libraries were quantified by qPCR and run on an Illumina NextSeq 500 (150bp single-end). Raw FastQ reads containing the amplicon primers (Supplemental Table 2, barcode_primers tab) were filtered (fastx-barcode-splitter, FastX-toolkit, http://hannonlab.cshl.edu) and used in subsequent analysis. The number of reads per DNA variant were quantified (fastX-collapser, FastX-toolkit) and the ChIP values were normalized to genomic DNA input. DNA variants were then sorted by the presence or absence of p53 or Sp1/KLF family motifs as determined by gimmeMotifs(27). Data tables for amplicon ChIP experiment can be found in Supplemental Table 4.

### Quantitative Real Time PCR

Total RNA was isolated using the Omega E.Z RNA kit with an on-column treatment with 50U of DNase I for 30 minutes. Single-stranded cDNA was generated with the High Capacity cDNA Reverse Transcription reagents (Applied Biosystems), and qPCR was performed on an Applied Biosystems 7900H with the relative standard curve method and iTaq Universal SYBR Green Supermix reagents (Biorad). qPCR primers are shown in Supplemental Table 2 (qPCR tab).

### Luciferase Plasmid Cloning and Expression Assays

Candidate enhancer sequences were synthesized as dsDNA (IDT) and cloned into the pGL4.24 destabilized Luciferase reporter vector (Promega) using the HiFi DNA Assembly method (NEB). Enhancer variants were created using the inverse PCR method with Hot Start Q5 Polymerase (NEB) and primers available in Table 2. Plasmids were reverse transfected according to manufacturer’s recommendations (JetPrime, Polyplus Transfection) in triplicate in a 96 well plate. Plasmid DNA (0.2µg) was transfected at a ratio of 9:1 for the candidate enhancer:constitutive promoter driving Renilla luciferase (pGL4.75, Promega). Luciferase activity was determined using the Promega Dual-Luciferase® Reporter Assay System according to manufacturer specifications on a Synergy HI plate reader (Bio-Tek).

### Western blotting

Total protein was isolated using RIPA buffer, followed by a 15-minute incubation on ice, and pelleting of and centrifugation to remove insoluble debris. Protein lysate samples were run at 180v volts on NuPAGE 10% Bis-Tris protein gels from Invitrogen. Samples were blotted onto 0.2 µm nitrocellulose membrane and probed with p53 (clone DO1, BD Biosciences, #554923) or GAPDH (Cell signaling #5174S) antibodies. Proteins were visualized with HRP-conjugated secondary antibodies and SuperSignal Chemiluminescent reagents (Thermo Scientific) and imaged on a BioRad Chemidoc imager.

### Protein expression and purification

Human p53_1-393_ in pGEX-2TK (Ampicillin) coding for an in-frame N-terminal GST tag was a gift from Cheryl Arrowsmith (Addgene plasmid # 24860; http://n2t.net/addgene:24860; RRID:Addgene_24860 and (33)). p53_1-393_ was expressed in Rosetta 2 DE3 cells (EMDMillipore) in 2YT media. Cells were grown at 37°C with shaking at 225rpm until OD_600nm_ = 0.4-0.6, then shifted to 22°C and induced with 1mM IPTG for 4 hours. Cells were harvested by centrifugation at 4800rpm for 15 minutes. Cell pellets were lysed in GST Buffer A (20mM TrisHCl pH7.3 (RT), 300mM NaCl, 0.2mM EDTA, 1mM DTT, 5% Glycerol, 10mM Na-Butyrate and supplemented with Complete protease inhibitor cocktail (Roche) and 0.1% NP40. Cells were lysed by sonication. Cell lysates were cleared by centrifugation at 18000 rpm for 45 minutes at 4C. Cell lysates were filtered with a 0.45uM syringe filter and loaded on to an equilibrated 5ml GSTrap HP column (GE Healthcare) using an Akta Purifier FPLC (GE Healthcare). The column was washed with GST Buffer A; bound protein was eluted using a linear gradient of 0-100% GST Buffer B (GST Buffer A + 10mM Glutathione (reduced)). Purity was determined by SDS PAGE using a 4-12% Bis-Tris gel run in 1x MOPS SDS buffer. Eluted fractions containing p53_1-393_ constructs were pooled and concentrated using a Vivaspin 10KDa MWCO centrifugal concentrator (GE healthcare). Concentrated protein was dialyzed overnight into GST-Buffer D (GST Buffer A + 20% glycerol). Protein concentration was determined using a nanodrop with calculated extinction co-efficient and molecular weight parameters. Purified protein was divided into single use aliquots, flash frozen in liquid nitrogen and stored at −80C.

### CCNG1 p53 electrophoretic mobility shift assay

Binding of p53 to CCNG1 enhancer sequences was tested by electrophoretic mobility shift assays (EMSAs). 60bp HPLC purified DNA sequences (IDT, Supplemental Table 2, EMSA tab) were resuspended at 100µM in 1x TE buffer. Equal volumes of complementary oligonucleotides were heated at 95°C for 5 minutes and annealed by cooling to 21C for 30 minutes. Binding reactions were assembled in DNase/RNase free, low adhesion microcentrifuge tubes. Reactions contained 1x p53 binding buffer (50mM Tris HCl pH 7.3, 150mM NaCl, 2mM DTT, 0.1mg/ml BSA). Unless otherwise stated, 50nM of annealed DNA was added to each reaction. Binding reactions were started by adding the required concentration of p53_1-393_ and allowed to proceed for 30 minutes at 21°C. Reactions were loaded immediately on a vertical 0.7% Agarose gel (Ultrapure agarose, Life Technologies) buffered with 1X TBE. Gels were run for 45 minutes at 60V in 1X TBE buffer. Gels were stained for 10 minutes with 1x SYBR gold nucleic acid stain (ThermoFisher) in 1x TBE buffer and imaged on a G:box blue light transilluminator (Syngene). Densitometric analysis was carried out in Fiji (34); binding curves were fitted using the Hill equation with a 2-site binding model (Hill co-efficient = 2) using non-linear regression (nls) in R (31).

### RNA sequencing

HCT116 parental, TP53-/-, or ATF3-/- cells were treated at 80% confluency in a 6-well plate with either DMSO or 5µM Nutlin-3A for 6 hours and total RNA was isolated (EZ RNA, Omega Biotek). PolyA+ RNA was purified using poly dT magnetic beads (Perkin Elmer) and fragmented at 90°C for 15 minutes. Fragmented RNA was used as the template for double-stranded cDNA production using random hexamers (1^st^ strand synthesis) and the dUTP method to preserve strandedness (2^nd^ strand synthesis). The resulting double-stranded cDNA was then used to construct an Illumina-compatible sequencing library (BioO NextFlex RNA Library Kit, Perkin Elmer). Libraries were quantified using qPCR (NEBNext Library Quantification, New England Biolabs) and an Agilent Bioanalyzer and then pooled for sequencing on an Illumina NextSeq 500. Sequencing reads were aligned to the hg19/GRCh37 assembly and transcript counts were determined using quantMode in STAR (35). Differential gene expression and normalized gene counts were determined using DESeq2 (30).

### Transcription factor peak intersections

Transcription factor peak files were obtained from the Cistrome database ((36), rycistrome.org/db, accessed July 1^st^, 2019). MPRA regions were converted into the hg38 genome assembly coordinates using liftOver from hg19 to hg38 (UCSC). Transcription factor peak summits were then intersected with MPRA regions using BedTools (intersectBed)(37). Peak intersection data were clustered by row (MPRA regions) and column (transcription factor) by One minus Pearson Correlation with complete linkage using Morpheus (https://software.broadinstitute.org/morpheus). Data underlying the cluster analysis can be found in Supplemental Table 5.

### Transcription factor motif analysis

Three tools were used to identify motifs either present or enriched within the 100bp regions from the MPRA p53-bound and p53-unbound peaks. Additionally, we analyzed transcription factor motif presence and enrichment within a set of p53 binding sites identified across multiple cell types in Verfaillie et al (9). For the Verfaillie p53 set, we considered only the 1001/1149 (87%) p53 peaks found within DNAse Hypersensitive Clusters (UCSC GenomeBrowser, ENCODE Regulation Track (38, 39)). Motif analysis was then performed on the entire DNAse Hypersensitive Cluster region. Known Motif enrichment using HOMER (40) was performed using the *findMotifsGenome* module (findMotifsGenome.pl -nomotif -size given) against the hg19 genome. Analysis for the presence of known motifs within MPRA regions and the Verfaillie p53 set was performed using the 2018 release of JASPAR Vertebrate Transcription Factor Motif Database (41) and both gimmeMotifs (27) and UCSC TableBrowser (42). Analysis of transcription factor motif enrichment using gimmeMotifs was run using the *scan* module (options -t -g hg19 -p JASPAR2018_vertebrates.pfm). The presence of hg19-based JASPAR vertebrate transcription factor motifs within p53-bound MPRA peaks and within the Verfaillie p53 set was performed using the UCSC TableBrowser and the JASPAR 2018 hg19 Track Hub. Only motifs with an enrichment score of 400 or higher (p < 0.0001) were considered for analysis. Histograms of motif enrichment within MPRA p53 or Verfaillie p53 peak sets were generated using bedTools (coverageBed -d option) and Morpheus.

### Sequencing data availability

All sequencing data generated as part of this manuscript is currently being submitted to Gene Expression Omnibus (GEO) and will be available shortly. p53 ChIP data from Nutlin-3A-treated cells were obtained from GSE86222. Conserved element data (bigWig) from PhyloP (43) and PhastCon (44) were obtained from the UCSC Genome Browser and plots were generated using deepTools (45).

## Results

We designed a barcoded and chromosomally integrated reporter system to assess the function of DNA sequences flanking the p53RE in p53-dependent CREs (Fig. 1A). The lentivirus-based system expresses eGFP under the control of a minimal promoter and a putative p53-dependent CRE sequence (29). Each putative CRE was included in the library with five unique, 12 nucleotide long barcodes encoded in the 3’UTR of eGFP to allow a molecular readout of transcriptional activity (Fig.1A). We selected 296 putative p53 binding locations based on the presence of a canonical p53 family binding motif, distance from the nearest transcriptional start site (< 100kb), and localization within a DNAse hypersensitive site (DHS). The 20bp p53 family motif was centered in a 100bp fragment with 40bp of flanking genomic context on each side. We also included 196 p53-independent and constitutively active CREs as determined by FANTOM Consortium CAGE data as positive controls of CRE activity (46). These sequences were cloned into the lentiviral plasmid backbone pLS-MP as previously described (29).

**Figure 1.**
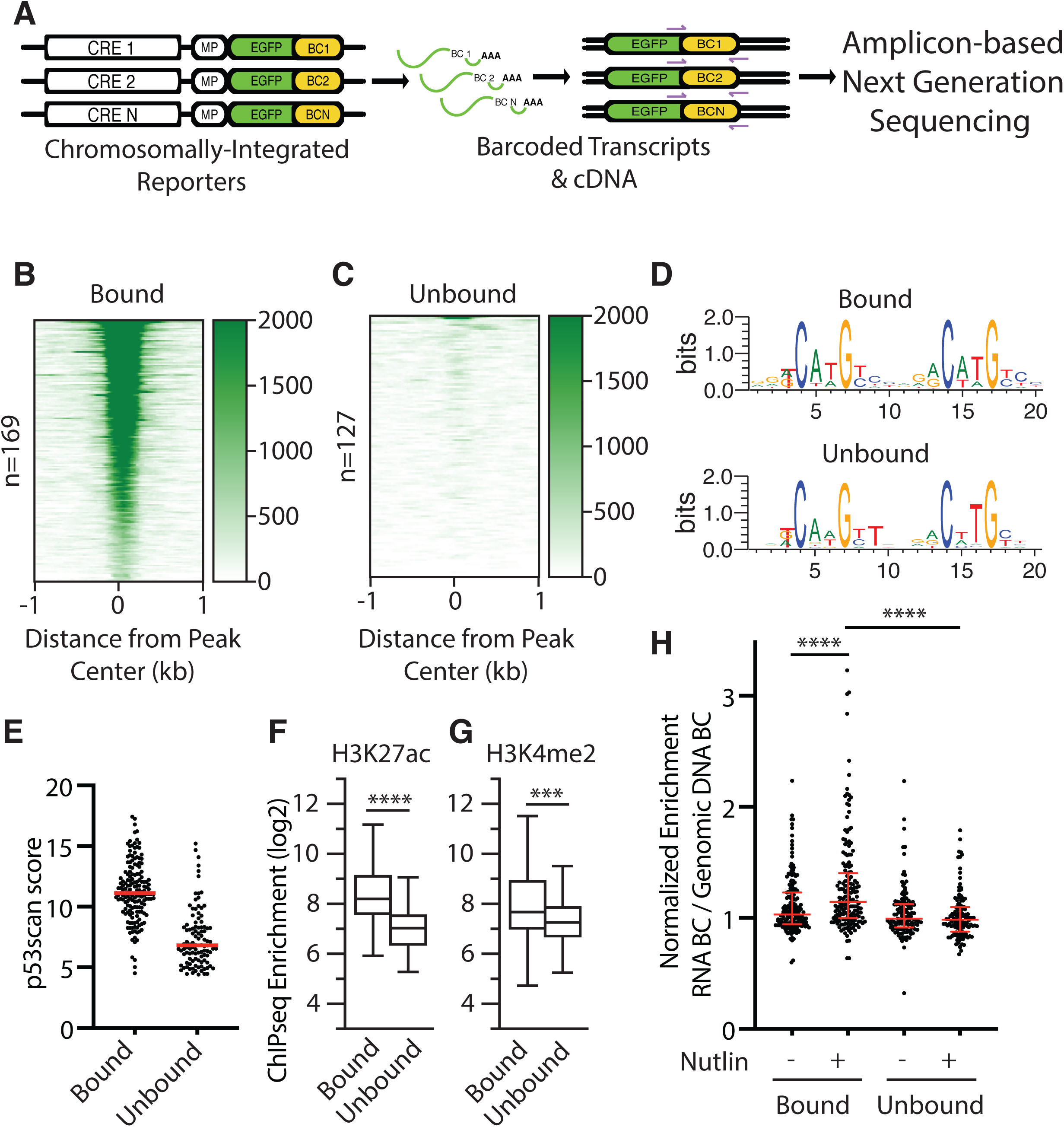
**(A)** Schematic and workflow for a massively parallel reporter assay (MPRA) to study the effect of flanking sequence context on p53 transcriptional activity. Heatmaps p53 ChIP-seq enrichment from Nutlin-3A-treated HCT116 colon carcinoma cell lines for **(B)** p53-bound or **(C)** p53-unbound regions found in the MPRA pool. **(D)** DNA sequence weight motifs for p53-bound (top) or unbound (bottom) regions from the MPRA pool. **(E)** Jitter plot of scores from p53scan (27) depicting the adherence to a canonical p53 family motif sequence for the p53-bound and p53-unbound regions. Enrichment of **(F)** H3K27ac or **(G)** H3K4me2 in Nutlin-3A-treated HCT116 cells at p53-bound or unbound regions. Statistics represent an unpaired, two-tailed T-test (*** p<0.001, **** p< 0.0001). **(H)** Normalized transcriptional activity of p53-bound or unbound regions from the MPRA assay after 6 hours of either DMSO or 10uM Nutlin-3A treatment. Data are depicted as enrichment of the 3’UTR-encoded barcode linked to each putative regulatory region normalized to the enrichment of that same sequence integrated into the genome and represent the averages from three biological replicates of each condition. (**** p<0.0001 by one-way ANOVA)

We then examined the activity of our putative CREs using the model human colon carcinoma cell line HCT116 which is well-suited for studying p53-dependent transcriptional activity. Our 296 potential p53-dependent CREs clustered into two groups based on p53 occupancy using ChIP-seq data from HCT116 cells (1). 169 out of 296 regions were scored as p53 binding sites (peaks) by MACS (Bound, Figure 1B), whereas 127 regions lacked measurable p53 binding (Unbound, Figure 1C). The average position weight matrix of the p53RE for each of the clusters is highly similar (Fig. 1D), however, the consensus motif in p53-bound regions more closely resembles the optimal p53 consensus motif than do p53-unbound CREs (Figure 1D-E). CREs bound by p53 show higher enrichment of canonical enhancer-associated histone modifications H3K27ac (Fig. 1F) and H3K4me2 (Fig. 1G) than do those regions lacking p53 occupancy in HCT116 cells.

We then performed triplicate biological measurements of p53-dependent CRE activity in HCT116 using our MPRA approach. MPRA-transduced cells were treated for 6 hours with either DMSO or the MDM2 inhibitor Nutlin-3A. Nutlin-3A leads to stabilization and activation of p53 at a similar level to what is seen with DNA damaging agents like etoposide (Fig.1H, 2A) (47). Importantly, enhancer activity measurements across biological replicates, treatment conditions, and cell lines were highly correlated (Supplemental Figure 1). As expected from our analysis of p53 occupancy, p53-bound CREs showed a bulk Nutlin-3A-dependent increase in activity compared to treatment with DMSO (Fig.1H, **** p<0.0001, one way ANOVA). Unbound CRE activity was unaffected by Nutlin-3A treatment relative to DMSO and was substantially lower than p53-bound CREs (Fig.1H, **** p<0.0001, one way ANOVA). Activity of the ubiquitously expressed CRE controls were unaffected by the induction of p53 by Nutlin-3A (Supplemental Fig. 2A). These results are consistent with *in vivo* p53 occupancy, p53 motif score, and previous observations that p53 binding correlates with increased transcriptional activity of CREs (2, 9, 10, 48).

**Figure 2.**
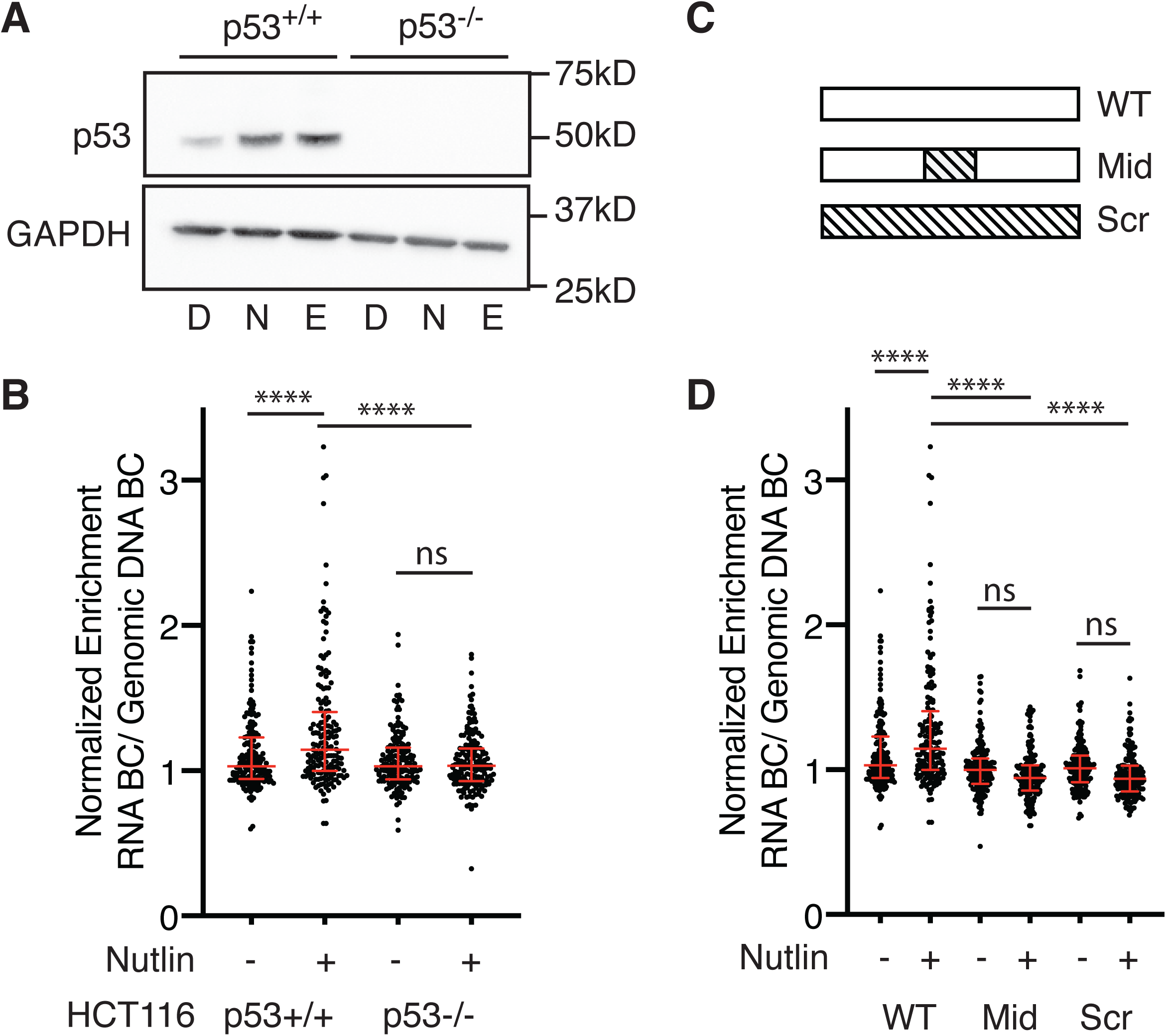
**(A)** Immunoblotting for p53 (top) or GAPDH (bottom) expression in HCT116 p53+/+ or p53-/- colon carcinoma cells after 6 hours of treatment with DMSO (D), 10uM Nutlin-3A (N), or 100uM etoposide (E). **(B)** Normalized transcriptional activity of the wild-type p53-bound regions in either HCT116 p53+/+ or p53-/- cells after a 6 hour treatment of either DMSO or 10uM Nutlin-3A. Data are depicted as enrichment of the 3’UTR-encoded barcode linked to each putative regulatory region normalized to the enrichment of that same sequence integrated into the genome and represent the averages from three biological replicates of each condition. (**** p<0.0001 using an ordinary one-way ANOVA). **(C)** Sequences within the wild-type p53-bound regions from the MPRA (WT) were shuffled (while preserving GC content) to alter either the 20bp p53 binding site (Mid) or the entire 100bp MPRA sequence. **(D)** Normalized transcriptional activity of p53-bound sequences for the wild-type (WT), p53-binding site scramble (Mid), or the full scramble (Scr) regions in HCT116 p53+/+ cells after a 6 hour treatment of either DMSO or 10uM Nutlin-3A. Data are depicted as enrichment of the 3’UTR-encoded barcode linked to each putative regulatory region normalized to the enrichment of that same sequence integrated into the genome and represent the averages from three biological replicates of each condition. (**** p<0.0001 by one-way ANOVA).

### p53-bound CREs require direct-binding of p53 for Nutlin-3A-induced activity

In order to determine if Nutlin-3A-induced activity of p53-bound CREs is p53-dependent, we assessed transcriptional activity in matched HCT116 TP53+/+ and TP53-/- cell lines (Fig.2A). Nutlin-3A-induced activity of p53-bound CREs was diminished in HCT116 TP53-/- cells suggesting these enhancers are dependent on wild-type p53 (Fig. 2B). As expected, ubiquitously expressed CRE controls were unaffected by the loss of p53 expression (Supplemental Fig. 2). To test whether CRE activity is direct or indirectly dependent on p53, we compared wild-type CRE sequences to those with either the 20bp p53RE (Mid) or the entire CRE sequence (Scr) randomized (Fig 2C). As a control for randomization, we fully scrambled the 196 ubiquitous CRE control sequences while preserving GC content, leading to a loss of activity (p<0.0001, one ANOVA, Supplemental Figure 2B). Scrambling either the p53RE or the entire CRE sequence abrogates Nutlin-3A-dependent enhancer activity (Fig. 2D, **** p<0.0001, one way ANOVA). In aggregate, wild-type CREs are more highly active than Mid or Scr CREs, suggesting that p53 strongly influences CRE function (Fig. 2D). Taken together, these data suggest that Nutlin-3A-induced activity of CREs requires direct binding of p53, in agreement with previous observations (9, 10).

### Variation in flanking sequence context alters p53-dependent CRE activity

Previous work suggests that p53-dependent CREs uniquely work in a single-factor mechanism in which only the presence of p53 is required for activation of transcription (9). Thus, this model implies that no other DNA binding factors are required for CRE activity. Similarly, analysis of DNA sequences flanking functional p53REs revealed no consistent enrichment or requirement for other transcription factor binding motifs or sequences outside of the p53RE (9, 10). Because this model represents a potential novel mechanism for enhancer function and diverges from canonical CRE models, we sought to directly test whether p53-dependent CRE activity requires sequences or transcription factor motifs outside of the p53RE. We systematically scrambled non-overlapping 20bp regions of each CRE starting at the 5’ end (Fig. 3A). As previously discussed, we also included controls where the p53RE and the entire CRE sequence were randomized. We then asked whether individual CRE variants had significantly different transcriptional activity than their wild-type counterpart in Nutlin-3A-treated, wild-type HCT116 cells. The large majority of Mid and Scr variants showed decreased activity relative to wild-type sequences, in line with p53-dependent *trans*-activation (Fig. 3D, one-way ANOVA with Tukey HSD, P value < 0.05). 93 Mid or Scr variants had statistically significant differential activity relative to the wild-type CRE, with 88 (95%) displaying reduced activity when sequences were randomized (Fig. 3D). Scrambling sequences flanking p53REs does not affect transcriptional activity in aggregate (Fig. 3B). However, many individual CREs have significantly altered transcriptional outputs when context-specific regions are scrambled (Fig.3C, one-way ANOVA with Tukey HSD, P value < 0.05). 48/83 (57%) p53RE-flanking variants have statistically reduced activity relative to the wild-type CRE, suggesting the original sequence positively regulates p53-dependent activation. Variants with decreased p53-dependent CRE activity had wild-type counterparts with higher expression values than compared to those variants with increased activity (Fig. 3E, p<0.0001, Mann-Whitney U). Our data demonstrate that changing sequences flanking the p53RE can alter p53-dependent CRE activity and suggests that additional DNA-encoded information beyond the p53RE is required for CRE function.

**Figure 3.**
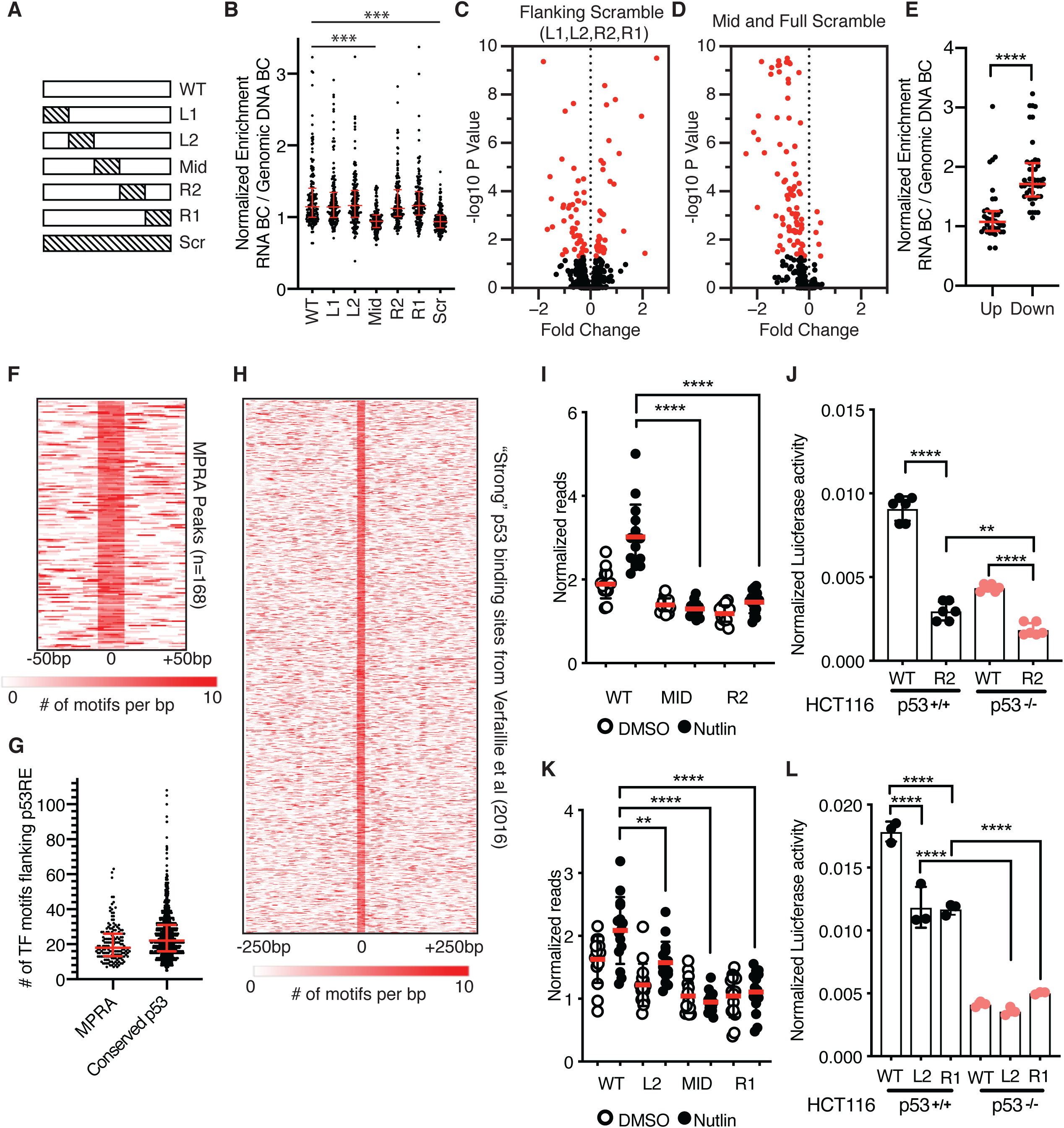
**(A)** Schematic depicting scrambling of 20bp sequences within 100bp p53-bound MPRA regions. Sequence scrambling was performed to preserve total GC content within the 20bp scrambled region relative to the wild-type sequence. **(B)** Normalized transcriptional activity of the MPRA sequences depicted in *(A)* in HCT116 p53+/+ cells after a 6 hour treatment with 10uM Nutlin-3A. Data are depicted as enrichment of the 3’UTR-encoded barcode linked to each putative regulatory region normalized to the enrichment of that same sequence integrated into the genome and represent the averages from three biological replicates of each condition. (*** p<0.001 using an ordinary one-way ANOVA). **(C)** Volcano plot of wild-type MPRA sequence activity compared to flanking scramble (L1, L2, R2, or R1). Data are plotted as Fold Change (WT over scramble) versus -log 10 *p-*value (from a one-way *ANOVA* with Tukey post-hoc test). **(D)** Volcano plot of wild-type MPRA sequence activity compared to Mid or Full scramble. Data are plotted as Fold Change (WT over scramble) versus -log 10 *p-*value (from a one-way *ANOVA* with Tukey post-hoc test). **(E)** Jitter plot depicting CRE activity values for wild-type p53-bound CREs (in the Nutlin-3A-treated condition) for those variants with increased (up) or decreased (down) activity relative to wild-type. P-value represents the result of an unpaired Mann-Whitney U Test (****, p<0.0001). **(F)** Heatmap of transcription factor motif enrichment across 100bp of the 168 p53-bound MPRA regions in the current study. Data represent a per base pair score for the presence of JASPAR-derived transcription factor motifs (p < 0.0001, corresponding to a JASPAR score of 400 or greater). Regions are centered on the putative p53 family response element. **(G)** Jitter plot displaying the number of JASPAR-derived transcription factor motifs found in a 100bp window (centered on a p53RE) for the 168 p53-bound MPRA regions or 1, 048 conserved p53 binding sites (taken from Verfaillie et al (9)). Median values and the interquartile range are depicted in red. **(H)** Heatmap of transcription factor motif enrichment across 500bp of the 1, 048 “strong” p53 binding sites from Verfaillie et al (9). Data represent a per base pair score for the presence of JASPAR-derived transcription factor motifs (p < 0.0001, corresponding to a JASPAR score of 400 or greater). Regions are centered on the putative p53 family response element and are extended −250bp and +250bp upstream. **(I)** Normalized transcriptional activity of the WT, Mid, or R2 version of Region 36/CCNG1 from the MPRA in HCT116 p53+/+ cells after 6 hours of either DMSO or 10uM Nutlin-3A treatment. (**** p<0.0001 using an ordinary one-way ANOVA). **(J)** Normalized Luciferase activity of either wild-type or R2-scrambled Region 36/CCNG1 sequence cloned upstream of a minimal promoter and driving expression of firefly luciferase in either HCT116 p53+/+ or p53-/- colon carcinoma cells. Firefly luciferase values were normalized to those of Renilla luciferase driven by a CMV promoter and co-transfected with the candidate Firefly plasmid (** p< 0.01, **** p<0.0001 using an ordinary one-way ANOVA) **(K)** Normalized transcriptional activity of the WT, L2, Mid, or R1 version of the TP53TG1 CRE from the MPRA in HCT116 p53+/+ cells after 6 hours of either DMSO or 10uM Nutlin-3A treatment. (** p<0.01, **** p<0.0001 using an ordinary one-way ANOVA). **(L)** Normalized Luciferase activity of either wild-type, L2, or R1-scrambled TP53TG1 CRE sequence cloned upstream of a minimal promoter and driving expression of firefly luciferase in either HCT116 p53+/+ or p53-/- colon carcinoma cells. Firefly luciferase values were normalized to those of Renilla luciferase driven by a CMV promoter and co-transfected with the candidate Firefly plasmid (**** p<0.0001 by one-way ANOVA)

To determine whether the flanking sequences might be disrupting additional DNA-encoded transcription factor binding information, we undertook a series of motif enrichment analyses. First, we examined the genome-wide enrichment of known transcription factor motifs across the p53-bound and p53-unbound MPRA regions using HOMER (40). Expectedly, p53 family motifs were highly enriched in the MPRA regions relative to size and GC-content matched genomic regions (Supplemental Table 6). We also observed statistically significant (Bonferroni q value < 0.05) enrichment of other known transcription factor motifs, including those in the AP-1, GATA, and ETS families (Supplemental Table 6). In order to account for potential bias in a single enrichment strategy, we used gimmeMotifs (27) and the JASPAR vertebrate database (41) to identify known transcription factor motifs found within our p53-bound MPRA regions. p53RE within MPRA regions are enriched for flanking transcription factor motifs (p-value < 0.0001, Fig. 3F-G), with a median of 18 distinct motifs found per p53-bound MPRA region. Examination of transcription factor ChIP data from the Cistrome Browser (36) suggests that many transcription factors can occupy p53-bound CREs (Supplemental Fig. 3). To ensure that the enrichment of transcription factor motifs was not an artifact of the regions we selected for our MPRA, we examined potential transcription factor motif enrichment within a validated set of p53 binding sites observed across multiple cell lines (9). Consistent with our p53-bound MPRA regions, we observe statistically significant enrichment of many transcription factor motifs flanking validated p53 binding sites using both HOMER and gimmeMotifs/JASPAR (Supplemental Table 8, Fig. 3G-H). This includes enrichment of AP-1, GATA, and ETS family motifs, but also an expansion of the types of enriched motifs, including SP1/KLF and Sox family motifs. Taken together, our data suggests that regions proximal to p53RE are enriched for transcription factor binding motifs that may be involved in p53-dependent CRE activity.

We then focused exclusively on CRE variants that displayed reduced activity relative to the wild-type CRE sequence. Our rationale was to identify potential transcription factors or functional DNA elements that are required for p53-dependent transcriptional activity. We first examined a CRE localized within the first intron of the *CCNG1* gene that is induced upon Nutlin-3A treatment in a p53-dependent manner (Fig. 3I). This activity is abrogated when either the p53RE or the 3’ adjacent 20bp region are scrambled (position R2, Fig. 3I). We observe similar results for a putative CRE localized within the second intron of *TP53TG1* where scrambling either the 5’ adjacent 20bp or 40bp downstream of the p53RE (Fig. 3K) leads to diminished p53-dependent CRE activity. We then sought to validate these MPRA results by utilizing a standard Luciferase reporter-based assay of CRE activity. In contrast to the 100bp sequence tested in the MPRA, we assessed the activity of a larger sequence encompassing an entire region of DNase hypersensitivity (DHS) as determined by ENCODE. DHS are putative regulatory regions often possessing transcriptional activity (38). Both the *CCNG1* and *TP53TG1* wild-type CREs are dependent on p53 for full activity (Fig. 3J, L, p53+/+), consistent with the MPRA data. We confirmed a reduction in CRE activity in HCT116 p53+/+ cells in both *CCNG1* and *TP53TG1* variants with reduced activity in the MPRA, suggesting our MPRA results are not an artifact of restricted sequence size (100bp) or assay conditions (Fig. 3J, L). For the *CCNG1* enhancer, loss of activity in the R2 variant is further reduced in the absence of p53 (Fig. 3J, p<0.01, one-way ANOVA), suggesting potential combinatorial activity of p53 and the wild-type R2 sequence within the *CCNG1* CRE. This combinatorial activity was not observed for *TP53TG1,* as the wild-type CRE and the L2 and R1 variants have similar activity in cells lacking p53 (Fig. 3L). These data suggest that p53-dependent CREs require different sequences and motifs flanking the p53RE for optimal activity.

### An SP1/KLF family motif is required for p53-dependent activity of the CCNG1 CRE

We continued investigating the role of flanking DNA sequences on p53-dependent CRE activity by further examining the *CCNG1* CRE. Both the DNA sequence within the R2 position and the p53RE are highly conserved across vertebrates suggesting that this region may have a conserved functional regulatory role (Fig.4A). The p53-dependence of the *CCNG1* CRE is similar across human cell types, as the R2 variant leads to a similar reduction in enhancer activity relative to the wild-type sequence when assayed in the non-transformed human cell line MCF10A (Fig. 4B). We then assessed the activity of the wild-type and R2 variant *CCNG1* CREs in *Trp53+/+* and *Trp53-/-* mouse embryonic fibroblasts (MEFs). The R2 variant has reduced activity compared to the wild-type *CCNG1* CRE in both wild-type and p53-deficient MEFs consistent with our observations in human cell lines (Fig.4C-D). In order to determine if the decrease in *CCNG1* CRE activity via R2 scrambling is due to a loss of the wild-type sequence or a gain of function from the scrambled sequence, we created a second R2 variant (R2*) that preserves GC content but is further randomized (Fig. 4A). Both R2 and R2* variants lead to loss of *CCNG1* CRE activity relative to wild-type and are further reduced in the absence of p53 (Fig. 4E). These results suggest that 20bp flanking the p53RE within *CCNG1* are required for p53-dependent CRE activity. The 20 bp immediately downstream of the p53RE within *CCNG1* enhancer contains several known transcription factor motifs based on analysis from JASPAR (41), suggesting that the loss of activity may be due to a loss of TF binding. The highest-scoring transcription factor motif from JASPAR are for members of the SP1/KLF family, and this G-rich motif is highly conserved across vertebrates (Fig.4F-G). We therefore made single base-pair mutations in the most conserved G3 and G4 residues of the SP1/KLF motif (Fig. 4F-G) and asked whether loss of these residues was sufficient to reduce CRE activity (49). Consistent with scrambling the entire 20bp R2 region, mutation of either the G3 or G4 position within the SP1/KLF motif severely diminishes transcriptional output in both wild-type and p53-deficient cell lines (Fig.4H-I). The additional reduction in CRE activity in p53-deficient cells seen in SP1/KLF motif variants suggests this region is functionally important independent of p53. Taken together, these data suggest that *CCNG1* CRE activity requires both p53 and a regulatory motif belonging to the SP1/KLF family.

**Figure 4.**
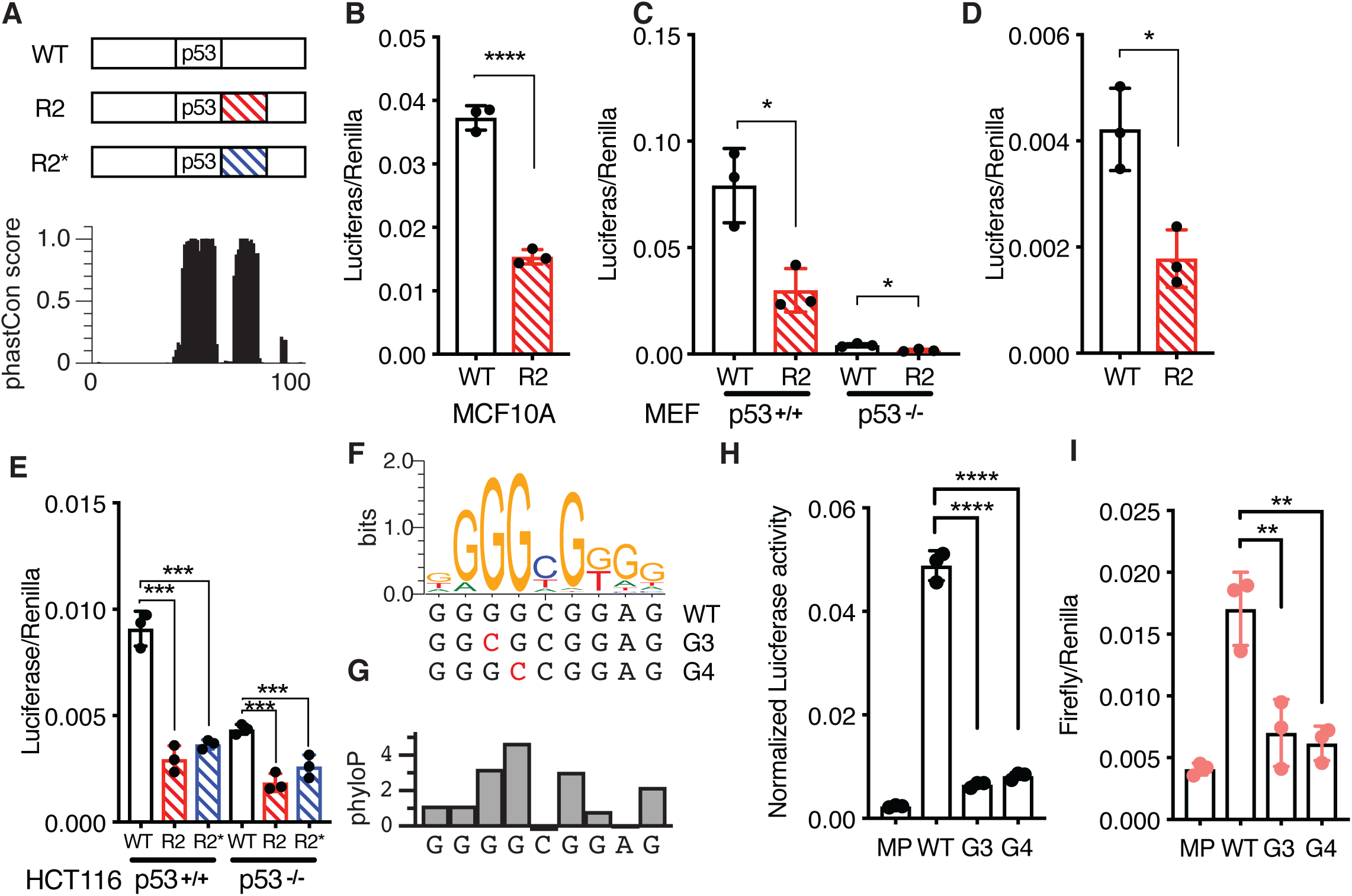
**(A)** Schematic of the 100bp Region 36/CCNG1 enhancer and per-basepair vertebrate phastCon score. **(B)** Normalized Luciferase activity (test sequence Firefly versus constitutive Renilla) for WT Region36/CCNG and the R2 version in the MCF10A mammary epithelial cell line. (**** p<0.0001 paired t-test) **(C)** Normalized Luciferase activity (test sequence Firefly versus constitutive Renilla) for WT Region36/CCNG and the R2 version in either p53+/+ or p53-/- mouse embryonic fibroblasts (* p<0.05, paired t-test). **(D)** Rescaled view of p53-/- reporter assay data from *(C)* depicting normalized Luciferase data for the wild-type or R2 version of the CCNG1 enhancer in p53-/- mouse embryonic fibroblasts. (* p<0.05, paired t-test) **(E)** Normalized luciferase activity for either the wild-type, R2, or R2* version of the CCNG1 enhancer in HCT116 p53+/+ or p53-/- cells. (*** p< 0.001 by one-way ANOVA). **(F)** The canonical Sp1/KLF family motif sequence in the CCNG1 enhancer as a transcription factor logo compared to the wild-type, G3, or G4 variants. **(G)** phyloP vertebrate conservation of the Sp1/KLF family motif within the wild-type CCNG1 enhancer sequence showing high conservation at the G3 and G4 positions. Normalized enhancer activity for the minimal promoter only (MP), WT, G2, or G3 CCNG1 variants in **(H)** HCT116 p53+/+ or **(I)** HCT116 p53-/- cells. (** p<0.01, **** p<0.0001, by one-way ANOVA)

### Loss of the SP1/KLF motif leads to reduced CCNG1 transcription and reduced p53 binding

Thus far, our data indicate that the p53-dependent *CCNG1* CRE also requires key DNA sequences flanking the p53RE and that these DNA sequences likely represents a binding motif for the SP1/KLF family. Therefore, we assessed the role of the p53RE and the R2 position sequence in their native genomic contexts using CRISPR/Cas9-mediated mutagenesis (Fig. 5A). Guide RNA sequences were targeted to either the p53RE, the R2 position, or the L2 position, which is 5’ adjacent to the p53RE, generating a population of indel mutations (Fig. 5A, Supplemental Table 4). Targeting of the p53RE substantially reduced endogenous *CCNG1* mRNA abundance relative to non-targeted Cas9 cells (Fig. 5B, p< 0.0001). Consistent with *in vitro* observations of CCNG1 enhancer activity, mutations within the R2 position reduce *CCNG1* mRNA levels, albeit not as severely as mutations in the p53RE (Fig. 5B, p<0.0001). As a control for sequence variants proximal to the p53RE, we generated indel mutations in the L2 position which is found in the 20bp immediately preceding the p53RE. Targeting the L2 position did not reduce endogenous *CCNG1* mRNA levels suggesting that proximal DNA mutations are not sufficient to decrease transcriptional output (Fig. 5B). Additionally, the L2 control suggests that the act of targeting Cas9 to the *CCNG1* intron does not affect *CCNG1* transcription on its own. These data further suggest that the p53RE and sequences immediately downstream of the p53RE are required for endogenous expression of the *CCNG1* mRNA.

**Figure 5.**
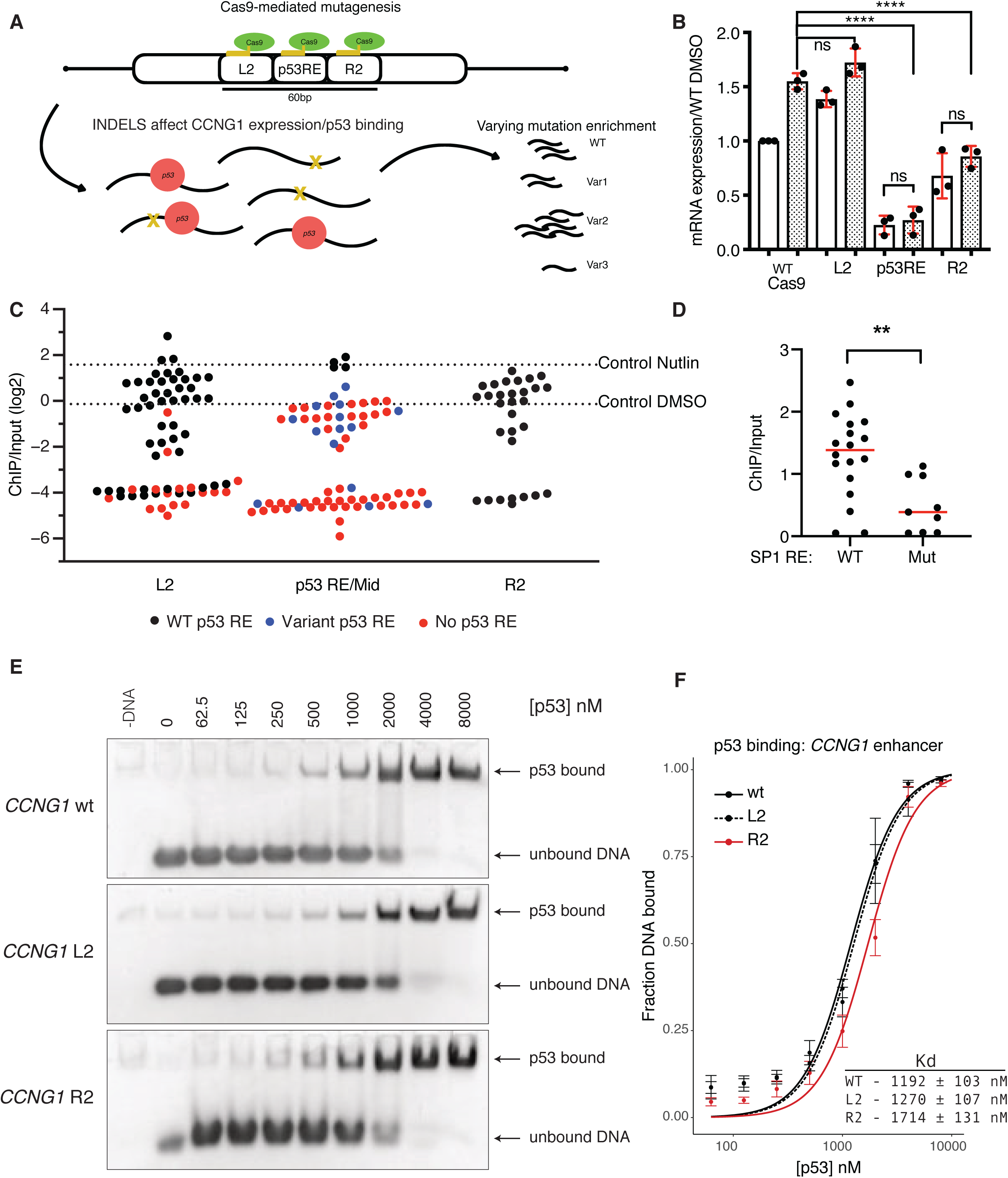
**(A)** Schematic depicting strategy to introduce insertion-deletion (indels) mutations into the native *CCNG1* locus in HCT116 cell lines. Pools of indel-containing cells were then used for measurement of *CCNG1* mRNA expression and for analysis of p53 binding by ChIP-sequencing. **(B)** qRT-PCR data of CCNG1 mRNA expression in control (WT Cas9), L2, p53RE, or R2-targeted Cas9 experiments after 6 hours of DMSO or 10uM Nutlin-3A treatment in HCT116 p53+/+ cell lines. (**** p<0.0001, by one-way ANOVA). **(C)** Jitter plot of p53 ChIP-enrichment for each Cas9-induced mutation at the native *CCNG1* enhancer. Black dots represent genetic variants that retain a wild-type p53 response element motif, while red dots are genetic variants with a mutated p53 response-element. Dashed line represents p53 ChIP/Input enrichment (log2) for the control cell line after 6hours of Nutlin-3A treatment. **(D)** Comparison of p53 ChIP enrichment from variants with either a canonical or mutated Sp1/KLF family motif from the R2-targeted Cas9 experiment. (red line is the median, ** p<0.01, unpaired t-test). **(E)** Electrophoretic Mobility Shift Assay (EMSA) for recombinant p53 bound to the wild type, L2, or R2 variant of the CCNG1 *cis-*regulatory element DNA sequence. Images are representative of 4 independent biological replicates. **(F)** Quantification of EMSA experiment in (E) performed as described in *Materials and Methods*. Fitted curves were fitted using a 2-site binding model (Hill equation; Hill co-efficient =2) using a non-linear regression.

We generated a pool of indel mutations at three locations within the *CCNG1* CRE, with mutations near the p53RE and the R2 position leading to a reduction in endogenous *CCNG1* mRNA expression (Fig. 5A-B). We therefore coupled chromatin immunoprecipitation of p53 to amplicon sequencing to simultaneously determine indel mutations and their potential effect on p53 occupancy at the *CCNG1* CRE. As a control, we performed p53 ChIP under DMSO and Nutlin-3A-treated conditions on HCT116 p53+/+ cells expressing wild-type Cas9 without a gRNA to target it to DNA. We then amplified a 150bp region in the *CCNG1* CRE from the p53-immunoprecipitated and input samples and sequenced them using Illumina approaches. We observed a 2.98-fold enrichment of the amplified sequence in the Nutlin-3A-induced samples relative to DMSO (Fig 5C, dotted lines), suggesting our amplicon ChIP approach was valid.

We next performed ChIP experiments from DMSO and Nutlin-3A-treated cell lines with Cas9 targeted to the p53RE, L2, or R2 positions. 66 unique DNA variants were identified within our pool when targeting Cas9 to the p53RE within the *CCNG1* CRE (Supplemental Table 4, Fig. 5C). As expected, the wild-type CCNG1 CRE sequence and 3 variants with an intact p53RE were enriched near the levels seen in the Cas9 control (Fig. 3C, p53RE, black dots). Enrichment of p53 is below or at DMSO levels when a p53RE is present but varies from the wild-type version (Fig. 5C, blue dots). As expected, *CCNG1* variants lacking a p53RE show a strong reduction in p53 binding. Variants proximal to the p53RE generally reduced p53 binding to the level of DMSO treatment (no enrichment), but many variants were depleted to the level seen when the p53RE was mutated (Fig.5C). These data are consistent with data showing that Cas9-induced mutations proximal to GATA1 binding sites alter GATA1 occupancy (50). None of the R2 mutations identified contained variants within the p53RE; however, when the conserved SP1/KLF motif was mutated or lost, p53 occupancy was reduced relative to the presence of an intact motif (Fig.5D, p<0.01). Our ChIP-based approach suggested that sequence variation flanking a p53RE can alter *in vivo* p53 binding in a context-dependent manner. We also performed electrophoretic mobility shift assays (EMSA) to better understand the effect of flanking sequence variation on p53 binding to p53RE. Increasing concentrations of recombinant p53 were combined with 60bp double-stranded DNA fragments representing the wild-type *CCNG1* p53RE and either the L2 or R2 variants from the original MPRA experiment (Fig. 5E). The R2 variant had a modest, but statistically significant, increase in K_d_ relative to either the wild-type or L2 variants (Fig. 5F). The reduction in p53 binding observed in the R2 variant by EMSA is consistent with our results from the *in vivo* variant ChIP experiment as well as the reduced p53-dependent transcription of endogenous *CCNG1.* Surprisingly, although a number of L2 variants had reduced p53 binding *in vivo,* we observed similar p53 binding affinities for both the wild-type and L2 sequences by EMSA (Fig. 5F). The fully randomized L2 variant does not affect p53-dependent CRE activity or affect p53 binding *in vitro,* suggesting that loss of p53 binding observed *in vivo* for specific L2 variants may be context-specific. These data, along with our examination of endogenous *CCNG1* mRNA expression, indicate that the specific sequences proximal to the p53RE in the *CCNG1* enhancer, which includes an SP1/KLF family motif, leads to increased p53 occupancy and higher *CCNG1* mRNA expression.

### p53-dependent transcription of GDF15 requires regulatory factors at two separate CREs

Data from our MPRA approach and follow-up experiments demonstrate that p53-dependent transcriptional activity at CREs is altered when sequences flanking the p53RE are perturbed. We then searched our dataset for additional p53-dependent regulatory regions with reduced activity upstream of known p53 dependent genes. One such putative region is approximately 11kb upstream of *GDF15,* which is well-characterized p53 target gene (2, 51). GDF15 has also recently been identified as a key modulator of the inflammatory and metabolic responses (52, 53). This region is enriched with the histone modifications H3K27ac and H3K4me2 and depleted for H3K4me3, a pattern strongly associated with transcriptional CREs. p53 is strongly bound to this region *in vivo* (Fig.6A). While examining the genomic context of this putative enhancer, we identified a second p53-bound, putative enhancer approximately 20kb upstream of *GDF15*. Both p53-bound regions are enriched for H3K27ac and H3K4me2 in the absence of p53 (Fig 6A) suggesting potential CRE activity independent of p53. This observation is consistent with recent reports of basal enhancer RNA transcription and histone modification enrichment at potential CREs in the absence of p53 (2, 48, 54). We therefore wanted to determine whether these p53-bound regions act as CREs for the endogenous expression of *GDF15*. We took advantage of a recently described approach to inactivate enhancers (55). In this approach, a catalytically inactive form of Cas9 (dCas9) was fused to the transcriptional repressor domain KRAB, and this strong repressor was targeted to either a control region (an enhancer for an unrelated gene, FGF2), the two p53-bound regions, or the *GDF15* promoter. Compared to the non-targeting control, targeting dCas9-KRAB to either p53-bound distal region (E1 or E2) reduced expression of endogenous GDF15 mRNA in both basal (DMSO) or p53-activated (Nutlin-3A) conditions (Fig. 6B). Repression of GDF15 mRNA levels when targeting E1 or E2 was similar to that of targeting the GDF15 promoter region (Fig. 6B). These results provide evidence that the E1 and E2 regions, bound by p53, are likely CREs regulating the expression of *GDF15*.

**Figure 6.**
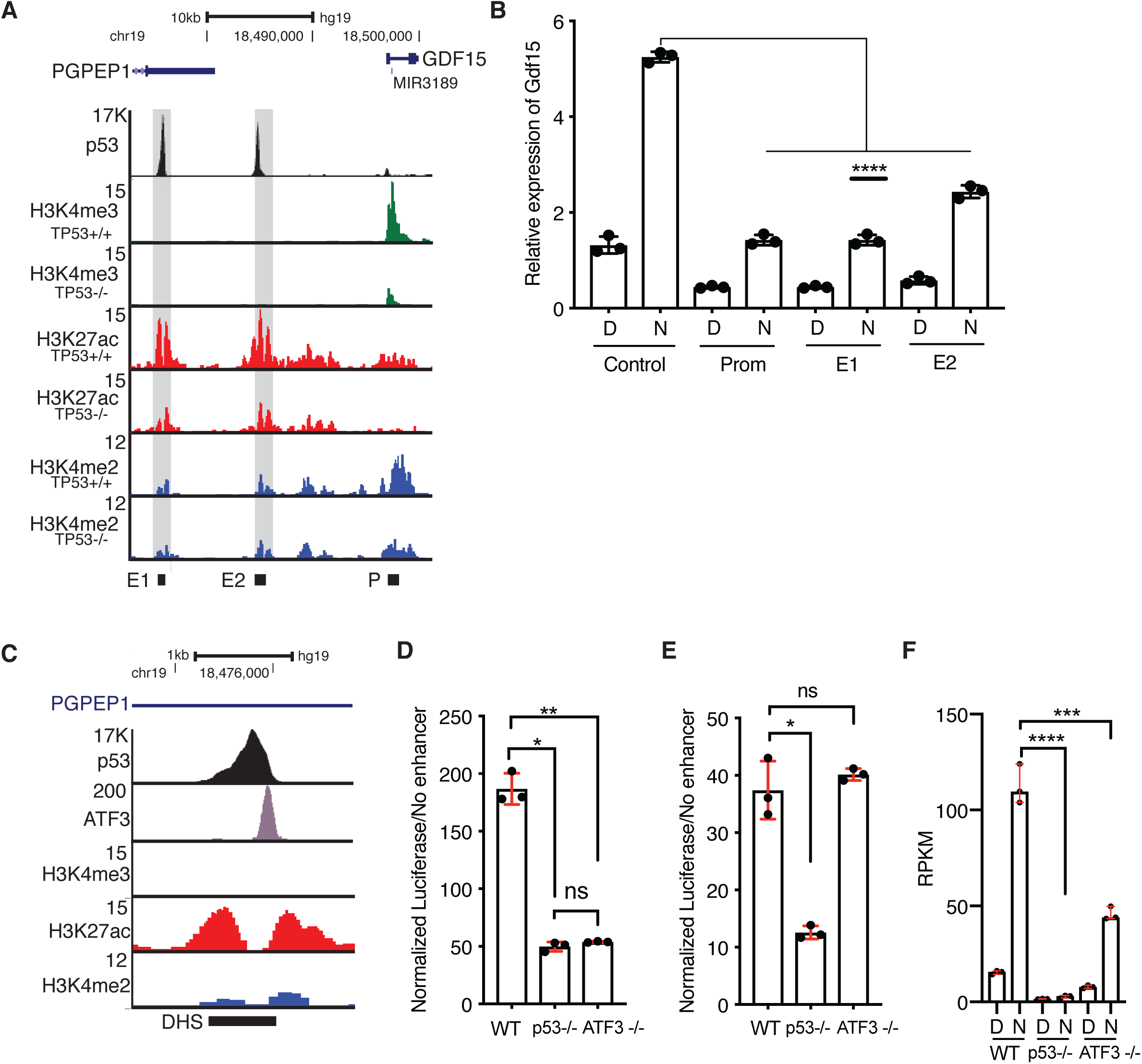
**(A)** Genome browser view of the *GDF15* locus showing p53, H3K4me3, H3K27ac, and H3K4me2 enrichment in HCT116 p53+/+ or HCT116 p53-/- cell lines. E1 = enhancer 1, E2 = enhancer 2, and P = *GDF15* promoter. Grey shaded boxes are placed over the E1 and E2 regions. **(B)** qRT-PCR relative expression of GDF15 mRNA in control, promoter, E1, or E2-targeted dCas-KRAB-expressing HCT116 p53+/+ cells treated with either DMSO (D) or 10uM Nutlin-3A (N) for 6 hours. (**** p<0.0001, one-way ANOVA). **(C)** Genome browser view of the GDF15 E1 region showing ChIP-seq enrichment of p53, ATF3, H3K4me3, H3K27ac, or H3K4me2. DHS = DNAse hypersensitive site. Normalized Luciferse activity (relative to minimal promoter only) of the GDF15 E1 enhancer **(D)** or the GDF15 E2 enhancer **(E)** in wild-type, p53-/-, or ATF3-/- HCT116 colon carcinoma cells. (* p<0.05, ** p<0.01, one-way ANOVA). **(F)** Reads per kilobase per million (RPKM) expression value of GDF15 mRNA from three replicates of polyA+ RNAseq of wild-type, p53-/-, or ATF3-/- HCT116 treated for 6 hours with either DMSO or 10uM Nutlin-3A. (*** p<0.001, **** p<0.0001, one-way ANOVA).

We next wanted to determine whether potential sequences or transcription factors might regulate the activity of these p53-bound CREs for *GDF15*. We sought to use the breadth of publicly available ChIP-seq datasets to identify potential transcription factors bound the GDF15 E1 CRE. Using information from the CISTROME database (36), we found that ATF3, a member of the AP-1 family of transcription factors, strongly binds to the GDF15 E1 CRE in HCT116 cells (Fig. 6C). The summit of the ATF3 binding event coincides with an ATF3 DNA motif, approximately 125bp downstream of the p53RE (Fig. 6C). We therefore focused on ATF3 because of its previous association as a positive regulator of p53 activity and its well-known role as a modulator of the inflammatory response (26, 56–58), of which GDF15 is a central regulator (53). Using a luciferase reporter approach, we assessed the activity of the GDF15 E1 enhancer in wild-type, p53-deficient, or ATF3-deficient HCT116 cells. Loss of either p53 or ATF3 leads to a substantial reduction of E1 enhancer activity (Fig. 6D). As expected by the lack of binding *in vivo,* the activity of the GDF15 E2 enhancer was unaffected by the loss of ATF3, whereas it is strongly dependent on p53 for activity (Fig.6D-E). Interestingly, although GDF15 E2 contains multiple AP-1 family motifs, it is not bound by or regulated by AP-1 member ATF3. Expression of endogenous GDF15 mRNA as determined by polyA+ RNAseq is reduced in either p53 or ATF3-deficient HCT116 cells consistent with *in vivo* binding and activity of ATF3 at the GDF15 E1 enhancer (Fig.6F). Taken together, our data indicate combinatorial activity of both p53 and ATF3 is required for activity of the GDF15 E1 enhancer and that both the E1 and E2 enhancer directly regulate expression of *GDF15*.

## Discussion

A number of high throughput analyses of p53 genomic occupancy revealed that p53 predominantly binds to *cis-*regulatory regions (CRE) like enhancers and promoters (34, 35, 39). The ability of p53 to activate transcription is well established, however, specific molecular mechanisms underlying p53 activity at CRE are less well studied. Here, we describe the use of a massively parallel reporter assay (MPRA) to characterize the effect of local sequence variation and transcription factor motifs on p53-dependent CRE activity. Our results support the canonical model of p53 activity where p53 primarily functions as a strong transcriptional activator at *cis*-regulatory elements (CRE) (61). Further, we confirmed that the p53 response element (p53RE) is a key predictor of p53-dependent transcriptional regulation as has been observed across multiple experimental systems (8–10). We then examined the contribution of local sequence context on enhancer activity by systematically altering sequences flanking the p53RE (Fig. 3A). Our results indicate that sequences outside of the core p53 binding site are required for optimal transcriptional activation (Fig. 3C). These functional sequences include both putative and confirmed transcription factor binding motifs, suggesting that p53 requires additional DNA-bound factors for its ability to activate transcription through CREs.

Two recent MPRA studies proposed a novel “single-factor” model for CRE regulation by p53 (9, 10). In this model, p53 is solely responsible for transcriptional output of an individual p53-bound CRE and does not require other transcription factors. Our data suggest that loss of p53 through either genetic depletion or through alteration of a p53RE sequence severely diminishes p53-dependent CRE activity (Figure 2B, D). Therefore, our data support the necessity of p53 for stimulus-dependent activation of p53-bound enhancers in agreement with the single-factor model (9, 10). Conversely, our results also demonstrate that additional transcription factors co-regulate p53-dependent CRE activity. Altering transcription factor binding sites proximal to the p53RE influences CRE activity to varying degrees. Our screening approach identified numerous motifs that positively or negatively affect CRE activity in a context-dependent manner (Fig. 3C). ATF3 binding to the GDF15 E1 enhancer is required for p53-dependent CRE activity and endogenous GDF15 mRNA expression (Fig. 6D-E). Additionally, a CRE regulating *CCNG1* transcription requires both a p53RE and an adjacent SP1/KLF family motif for activity (Figure 4E-F). Loss of either motif diminishes native *CCNG1* transcription driven by the CRE (Figure 5A). p53 occupancy at the *CCNG1* enhancer is reduced when sequences flanking the p53RE diverge from the wild type sequence as observed in our *in vivo* ChIP experiments (Fig. 5C). *In vitro* EMSA experiments suggest that only variation in the 3’ adjacent sequence, which contains the SP1/KLF motif, alters p53 binding affinity (Fig. 5E-F). Similarly, only mutations within the SP1/KLF motif affected endogenous *CCNG1* mRNA levels (Fig. 5B). CRE activity is reduced when evolutionarily conserved residues in the SP1/KLF motif are altered in both wild type and p53-deficient cells. These data suggest the *CCNG1* CRE requires both p53 and an additional factor bound to the SP1/KLF family motif for optimal activity.

The requirement for other transcription factors co-regulating p53-dependent transcriptional activity at CREs has not been characterized on a broad scale. Certainly, individual p53 CREs have been previously demonstrated to require co-regulatory factors in reporter assays, including the requirement for the p53RE and an AP-1 element bound by JunD for DNA damage-dependent activation of *GADD45A* transcription (62). A recent *in vivo* CRISPR/Cas9-screening approach identified CEBPB binding within a p53-dependent CRE required for optimal transcription of *CDKN1A* and initiation of senescence (22). Single nucleotide polymorphisms associated with lung cancer risk found within a p53-regulated CRE disrupt canonical transcription factor motifs, reduce p53 binding, and alter expression of *TNFRSF19 (7).* In this study, we identified both ATF3 and a likely member of the SP1/KLF family as co-regulatory transcription factors required for p53-dependent CRE activity. Of note, we have not identified specific transcription factors that bind to the *CCNG1* or *TP53TG1* CREs whose p53-dependent activity is altered upon variation in flanking sequence. In the case of *CCNG1,* the SP1/KLF family motif can be bound by over 12 family members, most of which are expressed in the cell type (HCT116 colon carcinoma) used in this study. Identification of specific transcription factors binding to individual DNA elements is often challenging, but updated approaches like the enhanced yeast 1-hybrid or quantitative mass spectrometry methods are now possible (63, 64). Importantly, we cannot rule out potential transcription factor-independent roles for these motifs in regulating p53-dependent CRE activity, such as the possibility that DNA shape or nucleosome positioning changes *in vivo* might be affected by changes in CRE sequence.

MPRAs are powerful tools for rapidly dissecting how sequence variation and context contributes to the activity of CREs. Despite their power, specific assay design choices and the non-native genomic context of the MPRA approach might help to explain some of the discrepancies between our work and previous reports (9, 10). First, we used a random, lentiviral-based genomic integration strategy to deliver our MPRA constructs, whereas previous p53-based approaches have used transient, plasmid-based delivery (9, 10). Genomic integration allows for the greater influence of chromatin and higher-order genomic structure which directly influence transcription factor binding and activity (29). In direct comparison with episomal DNA, the activity of integrated massively parallel reporter constructs was more reproducible and more closely aligned with expected activity based on CRE-associated biochemical patterns like histone modification and accessible chromatin in previous MPRA analyses (29). Second, our assay was specifically designed to test the effect of sequence variation on p53-dependent CRE activity while *post hoc* computational approaches were previously used to identify sequence features defining CRE activity. In both the primary MPRA screen and in traditional plasmid-based enhancer assays, we observed that variation in transcription factor motif sequences flanking the p53RE could alter CRE activity. Altering p53RE-adjacent TF motif sequences did not always alter CRE activity, suggesting sequence and context-dependent effects (Fig. 3C). Previous reports could not identify sequence-based features beyond the p53RE that predicted p53-bound CRE activity using machine learning and traditional motif enrichment approaches (9, 10). While accurate to say that there are no other transcription factor motifs or sequence-features that are as enriched as the p53RE (Supplemental Tables 6-7), we find that other transcription factor motifs are overrepresented in the CREs we studied and within a previously identified core set of p53-bound CREs (Supplemental Tables 6-7). This includes overrepresentation of the motif for the stress-dependent transcription factor ATF3 (Supplemental Table 6), which binds to a number of p53-dependent CREs and regulates activity of a CRE for *GDF15* (Fig. 6D-E). ATF3 is a well-studied regulator of p53-dependent transcription through control of p53 stability and co-factor recruitment (26, 56, 57). Previous reports clearly demonstrate that ATF3 can directly alter p53 stability and modulate p53 activity through interactions with histone modifying enzymes (56, 57, 65). Our work uniquely identifies a direct role for ATF3 DNA binding within a p53-bound CRE and demonstrates a positive effect on p53-dependent transcriptional activity. Given that ATF3 binds to numerous p53-bound regions of the genome (26) and their previously reported relationship, further examination into the local interplay between p53 and ATF3 at DNA is warranted.

p53 is a pioneer transcription factor and can mediate context-dependent chromatin remodeling at CREs (10, 48, 54). Despite this activity, the large majority of p53 genomic binding events occur within regions that are accessible before p53 engagement (10, 48, 54, 59), similar to what is observed for glucocorticoid receptor binding (66). These regions also contain chromatin modifications associated with active CRE, including H3K27ac and H3K4me1/2 before p53 binding (1, 48, 54, 59, 60, 60). Further, p53 depletion does not alter basal CRE-associated chromatin modifications or chromatin structure at the large majority of CRE (48, 54). Enhancer-derived RNA (eRNA) have been identified as a strong predictor of transcription factor binding and CRE activity (67). In further support of a multi-factor model, eRNA are transcribed from p53-regulated CREs in the absence of p53 (2, 48). We observe enrichment of enhancer-associated histone modifications H3K27ac and H3K4me2 at GDF15 E1 and E2 enhancers in the absence of p53 (Fig. 6A.) These data suggest that other factors are likely responsible for establishing and maintaining chromatin structure and basal activity at the majority of p53-bound CREs. Recent reports suggest that p53 binding and activity is strongly influenced by cell type-specific chromatin accessibility (1, 2, 54), which itself is controlled by differential transcription factor activity (11, 12, 68–70). How p53 functions across various cell and tissue contexts is still a vital and open question, but recent reports suggested that p53 binding and activity can be influenced by cell type (2, 54, 71). We propose that the lack of a core set of commonly enriched transcription factors within p53-dependent CREs is an important regulatory feature of the p53 network. By utilizing different sets of transcription factor co-regulators, we hypothesize that p53-dependent CRE activity is buffered against loss of any one regulatory partner. This hypothesis for p53 is strongly reminiscent of the “billboard” model seen at many *Drosophila* developmental enhancers (11, 18), whereby different combinations of factors can bind to a CRE and produce similar transcriptional outputs. The flexible billboard model for p53-bound enhancers is also consistent with a “distributed p53 network” whereby p53 transcriptionally controls many genes but that any one p53 target gene is dispensable for tumor suppression (1).

## Acknowledgments

We acknowledge the Center for Functional Genomics at the University at Albany for sequencing support. This work was supported by the National Institutes of Health NIGMS (R15 GM128049 to MAS) and startup funds from the University at Albany (to MAS). DB was supported by the Wellcome Trust (213501/Z/18/Z)

---

**Supplemental Figure 1.**
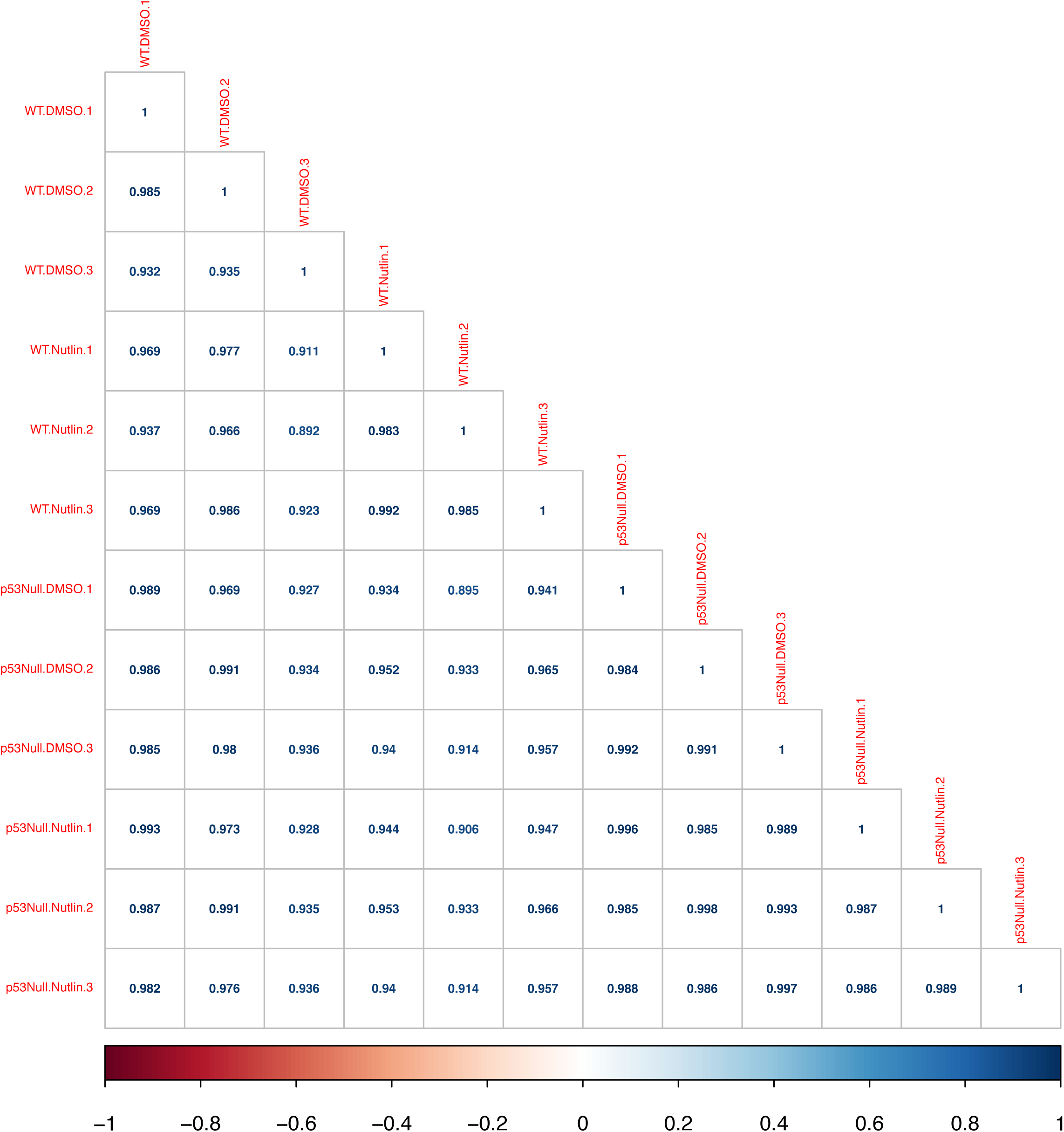
Pearson correlation values (R^2^) for raw enhancer barcode counts from three biological replicates across HCT116 p53+/+ and HCT116 p53-/- cells treated with either DMSO or 5uM of Nutlin-3A for 6 hours.

**Supplemental Figure 2.**
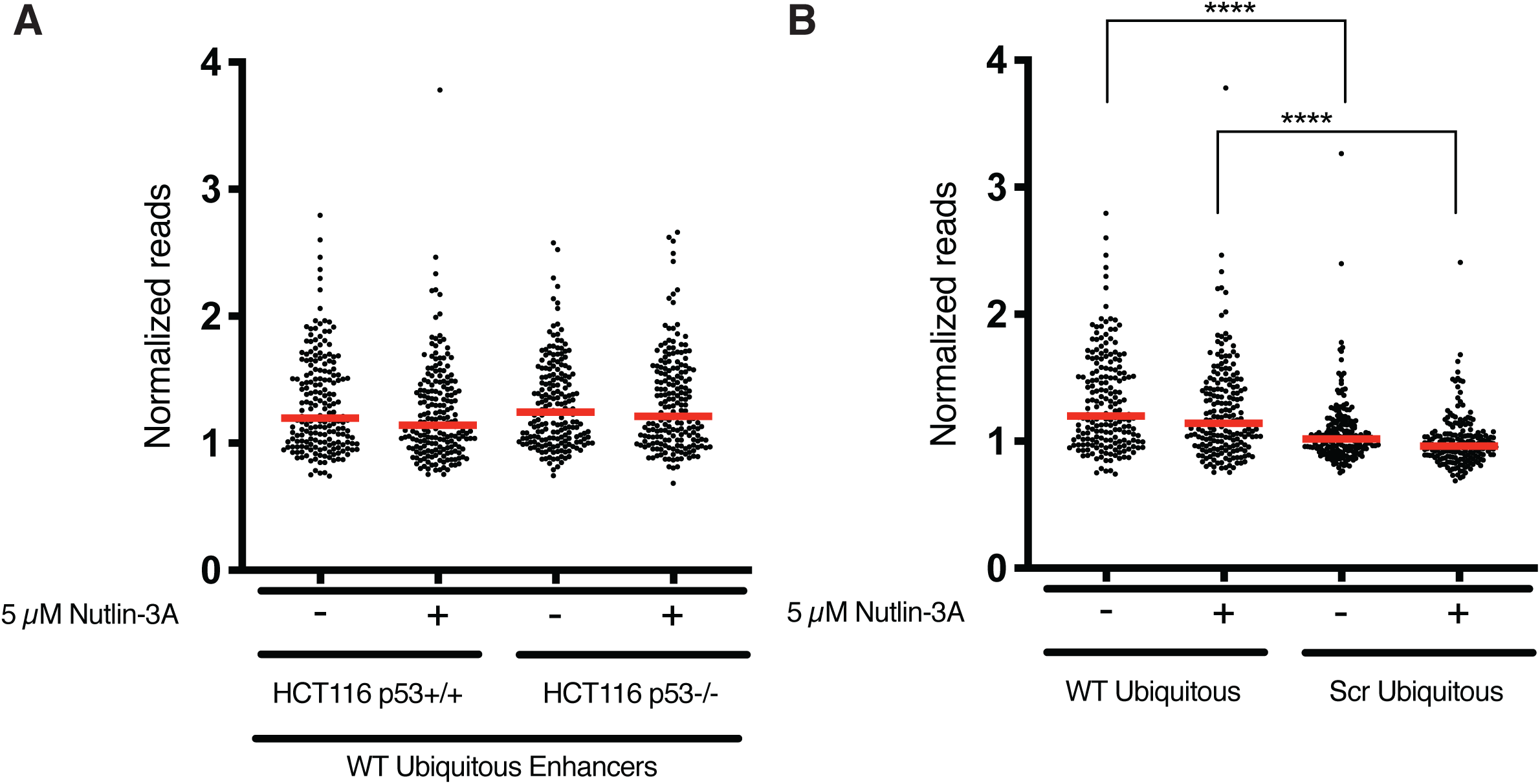
**(A)** Jitter plot of normalized CRE activity (RNA barcode/genomic DNA barcode) for 196 ubiquitously expressed CREs from the FANTOM Consortium in HCT116 p53+/+ and p53-/- cells treated with either DMSO or 5μM Nutlin-3A for 6 hours. Red line represents the median expression values. All conditions were statistically similar using a one-way ANOVA. **(B)** Jitter plot of normalized CRE activity (RNA barcode/genomic DNA barcode) for 196 ubiquitously expressed CREs or 196 randomized sequences from the FANTOM Consortium in HCT116 p53+/+ treated with either DMSO or 5μM Nutlin-3A for 6 hours (**** p<0.0001). Red line represents the median expression values.

**Supplemental Figure 3.**
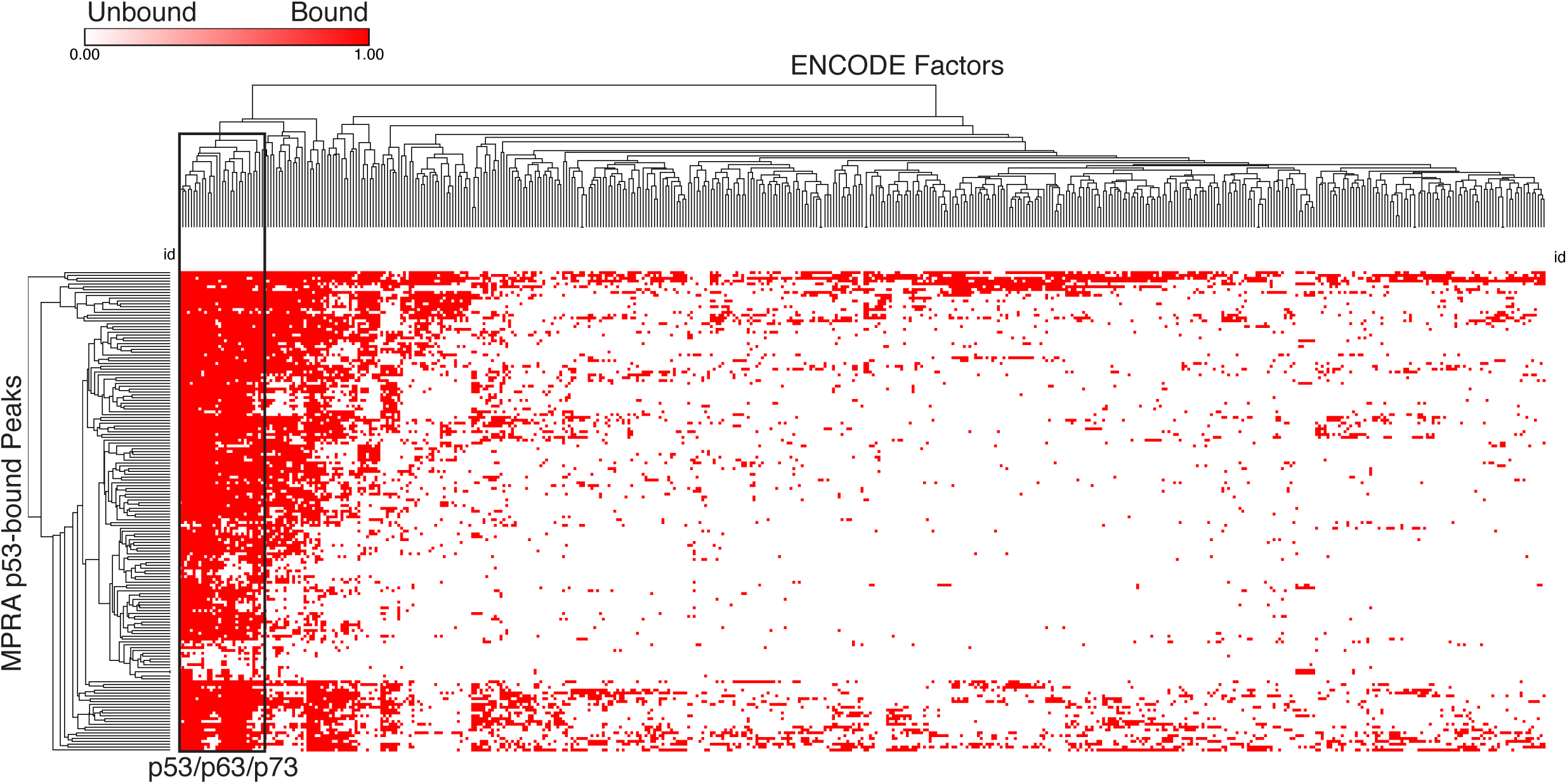
Hierarchical clustering of CISTROME-derived transcription factor binding with the 168 p53-bound MPRA regions from the current study. p53-bound MPRA regions are found on the Y-axis, while transcription factors bound to greater than 10 regions are found on the X axis. Only the peak summit for the CISTROM transcription factors were considered when intersecting with the p53-bound MPRA regions. Data were clustered by row and column using complete linkage and Pearson minus one correlation in the Morpheus software suite. The boxed region of the cladogram represents p53/p63/p73 datasets from the CISTROME database and shows strong enrichment of these factors with the MPRA regions from this study.

