## Supplemental Table 6 for "Locally acting transcription factors are required for p53-dependent *cis-*regulatory element activity"

| Name | Family | P-value | q-value (Benjamini) | % of Sequences w/ Motif | Fold Enrichment vs. Background |
|---|---|---|---|---|---|
| p53 | p53 | 0.00E+00 | 0 | 95.27 | 96.23 |
| p73 | p53 | 1.00E-281 | 0 | 81.66 | 157.04 |
| p63 | p53 | 1.00E-234 | 0 | 100 | 24.57 |
| ZNF416 | Zf | 1.00E-09 | 0 | 40.83 | 2.10 |
| Fra2 | bZIP | 1.00E-07 | 0 | 14.2 | 3.78 |
| Tcfcp2l1 | CP2 | 1.00E-06 | 0 | 7.69 | 6.52 |
| Atf3 | bZIP | 1.00E-06 | 0 | 15.98 | 2.93 |
| JunB | bZIP | 1.00E-06 | 0 | 14.2 | 3.13 |
| BATF | bZIP | 1.00E-05 | 0.0002 | 14.79 | 2.70 |
| Fosl2 | bZIP | 1.00E-05 | 0.0003 | 9.47 | 3.73 |
| AP-1 | bZIP | 1.00E-04 | 0.0004 | 15.98 | 2.47 |
| Fra1 | bZIP | 1.00E-04 | 0.0005 | 12.43 | 2.85 |
| ERG | ETS | 1.00E-04 | 0.0005 | 28.99 | 1.80 |
| TEAD4 | TEA | 1.00E-04 | 0.0011 | 18.93 | 2.11 |
| Gata4 | Zf | 1.00E-03 | 0.0033 | 17.75 | 2.04 |
| ETS1 | ETS | 1.00E-03 | 0.009 | 18.34 | 1.89 |
| ELF3 | ETS | 1.00E-03 | 0.0111 | 14.2 | 2.09 |
| FOXA1 | Forkhead | 1.00E-03 | 0.0128 | 2.96 | 7.40 |
| Nrf2 | bZIP | 1.00E-03 | 0.0144 | 2.96 | 7.22 |
| Zfp809 | Zf | 1.00E-03 | 0.016 | 6.51 | 3.16 |
| EWS | ETS | 1.00E-02 | 0.0211 | 13.61 | 2.00 |
| EHF | ETS | 1.00E-02 | 0.0403 | 20.71 | 1.63 |
| Gata6 | Zf | 1.00E-02 | 0.0444 | 14.2 | 1.85 |
