## Supplemental Table 7 for "Locally acting transcription factors are required for p53-dependent *cis-*regulatory element activity"

| Name | Family | P-value | q-value (Benjamini) | % of Sequences w/ Motif | Fold Enrichment vs. Background |
|---|---|---|---|---|---|
| p73 | p53 | 1e-1380 | 0 | 68.83 | 167.88 |
| p53 | p53 | 1e-1218 | 0 | 73.03 | 74.52 |
| p53 | p53 | 1e-1218 | 0 | 73.03 | 74.52 |
| p63 | p53 | 1e-790 | 0 | 72.33 | 19.82 |
| p53 | p53 | 1e-432 | 0 | 26.37 | 101.42 |
| Zfp809 | Zf | 1.00E-66 | 0 | 13.29 | 6.96 |
| ZNF416 | Zf | 1.00E-63 | 0 | 41.76 | 2.25 |
| Tcfcp2l1 | CP2 | 1.00E-45 | 0 | 8.19 | 8.03 |
| Smad4 | MAD | 1.00E-40 | 0 | 33.87 | 2.07 |
| Smad3 | MAD | 1.00E-29 | 0 | 47.25 | 1.56 |
| Smad2 | MAD | 1.00E-28 | 0 | 29.67 | 1.90 |
| TEAD4 | TEA | 1.00E-25 | 0 | 19.68 | 2.22 |
| Gata1 | Zf | 1.00E-24 | 0 | 13.49 | 2.70 |
| CArG | MADS | 1.00E-21 | 0 | 8.79 | 3.32 |
| Gata2 | Zf | 1.00E-21 | 0 | 13.79 | 2.45 |
| Gata6 | Zf | 1.00E-18 | 0 | 16.38 | 2.11 |
| NRF | NRF | 1.00E-15 | 0 | 3.90 | 5.13 |
| Gata4 | Zf | 1.00E-12 | 0 | 16.08 | 1.81 |
| BATF | bZIP | 1.00E-11 | 0 | 11.39 | 2.03 |
| TEAD | TEA | 1.00E-11 | 0 | 13.09 | 1.87 |
| GATA3 | Zf | 1.00E-11 | 0 | 22.18 | 1.56 |
| PRDM10 | Zf | 1.00E-10 | 0 | 12.89 | 1.84 |
| NPAS | bHLH | 1.00E-10 | 0 | 26.37 | 1.46 |
| TEAD1 | TEAD | 1.00E-10 | 0 | 16.48 | 1.67 |
| Fra2 | bZIP | 1.00E-10 | 0 | 8.39 | 2.16 |
| BMAL1 | bHLH | 1.00E-09 | 0 | 29.27 | 1.41 |
| Fra1 | bZIP | 1.00E-09 | 0 | 9.39 | 2.04 |
| AP-1 | bZIP | 1.00E-09 | 0 | 12.19 | 1.83 |
| KLF3 | Zf | 1.00E-09 | 0 | 8.39 | 2.11 |
| Fosl2 | bZIP | 1.00E-09 | 0 | 6.39 | 2.40 |
| Sp5 | Zf | 1.00E-09 | 0 | 13.19 | 1.76 |
| Atf3 | bZIP | 1.00E-09 | 0 | 10.79 | 1.89 |
| MNT | bHLH | 1.00E-09 | 0 | 18.38 | 1.57 |
| Jun-AP1 | bZIP | 1.00E-08 | 0 | 5.09 | 2.62 |
| Sp2 | Zf | 1.00E-08 | 0 | 19.78 | 1.51 |
| KLF5 | Zf | 1.00E-08 | 0 | 17.68 | 1.55 |
| TEAD3 | TEA | 1.00E-08 | 0 | 17.88 | 1.54 |
| JunB | bZIP | 1.00E-08 | 0 | 8.99 | 1.93 |
| CLOCK | bHLH | 1.00E-07 | 0 | 8.99 | 1.88 |
| Sox2 | HMG | 1.00E-07 | 0 | 14.19 | 1.61 |
| Sox10 | HMG | 1.00E-06 | 0 | 23.98 | 1.37 |
| SPDEF | ETS | 1.00E-06 | 0 | 13.99 | 1.51 |

| | | | | | |
|---|---|---|---|---|---|
| Bach2 | bZIP | 1.00E-05 | 0 | 3.80 | 2.42 |
| Sox3 | HMG | 1.00E-05 | 0 | 24.48 | 1.32 |
| FOXK2 | Forkhead | 1.00E-05 | 0 | 9.69 | 1.63 |
| Sox6 | HMG | 1.00E-05 | 0 | 22.08 | 1.34 |
| Sox4 | HMG | 1.00E-05 | 0 | 12.99 | 1.48 |
| Ets1-distal | ETS | 1.00E-05 | 0.0001 | 5.19 | 1.94 |
| PU.1 | ETS | 1.00E-05 | 0.0001 | 7.89 | 1.67 |
| NFAT | RHD | 1.00E-04 | 0.0001 | 12.39 | 1.47 |
| GATA:SCL | Zf,bHLH | 1.00E-04 | 0.0001 | 2.50 | 2.66 |
| NFkB-p65 | RHD | 1.00E-04 | 0.0003 | 7.99 | 1.59 |
| Ap4 | bHLH | 1.00E-04 | 0.0003 | 16.58 | 1.35 |
| CEBP:CEBP | bZIP | 1.00E-04 | 0.0004 | 3.50 | 2.11 |
| Six2 | Homeobox | 1.00E-04 | 0.0005 | 12.19 | 1.42 |
| Foxa2 | Forkhead | 1.00E-04 | 0.0007 | 10.89 | 1.44 |
| Foxf1 | Forkhead | 1.00E-03 | 0.0008 | 11.79 | 1.41 |
| NPAS2 | bHLH | 1.00E-03 | 0.0009 | 16.98 | 1.31 |
| Egr2 | Zf | 1.00E-03 | 0.0017 | 1.90 | 2.57 |
| ETV1 | ETS | 1.00E-03 | 0.002 | 16.78 | 1.30 |
| HLF | bZIP | 1.00E-03 | 0.002 | 11.29 | 1.39 |
| GABPA | ETS | 1.00E-03 | 0.0023 | 11.19 | 1.38 |
| Sox17 | HMG | 1.00E-03 | 0.0026 | 10.49 | 1.40 |
| Six1 | Homeobox | 1.00E-03 | 0.0029 | 3.80 | 1.82 |
| Etv2 | ETS | 1.00E-03 | 0.0031 | 12.19 | 1.35 |
| GATA3 | Zf | 1.00E-03 | 0.0032 | 1.80 | 2.50 |
| SpiB | ETS | 1.00E-03 | 0.0041 | 3.90 | 1.76 |
| FOXP1 | Forkhead | 1.00E-03 | 0.0044 | 5.79 | 1.56 |
| ZBTB18 | Zf | 1.00E-03 | 0.0046 | 7.89 | 1.45 |
| Foxo3 | Forkhead | 1.00E-03 | 0.0046 | 9.59 | 1.39 |
| FOXK1 | Forkhead | 1.00E-02 | 0.0064 | 12.69 | 1.31 |
| NFY | CCAAT | 1.00E-02 | 0.0083 | 9.49 | 1.37 |
| FoxL2 | Forkhead | 1.00E-02 | 0.0096 | 10.19 | 1.34 |
| Sox15 | HMG | 1.00E-02 | 0.0107 | 14.59 | 1.27 |
| Bach1 | bZIP | 1.00E-02 | 0.0164 | 1.10 | 2.75 |
| ELF3 | ETS | 1.00E-02 | 0.0173 | 9.29 | 1.33 |
| Tcf21 | bHLH | 1.00E-02 | 0.0235 | 12.29 | 1.27 |
| FOXM1 | Forkhead | 1.00E-02 | 0.0242 | 12.59 | 1.26 |
| ZNF467 | Zf | 1.00E-02 | 0.0272 | 10.39 | 1.29 |
| ZNF7 | Zf | 1.00E-02 | 0.0294 | 7.39 | 1.36 |
| TEAD2 | TEA | 1.00E-02 | 0.0301 | 7.69 | 1.34 |
| HIF-1a | bHLH | 1.00E-02 | 0.0348 | 3.00 | 1.64 |
| WS:ERG-fusio | ETS | 1.00E-02 | 0.0348 | 9.29 | 1.30 |
| ETS1 | ETS | 1.00E-02 | 0.0348 | 12.09 | 1.25 |
| NF1:FOXA1 | CTF,Forkhead | 1.00E-02 | 0.0348 | 1.00 | 2.56 |
| Elf4 | ETS | 1.00E-02 | 0.0348 | 11.79 | 1.25 |
| Maz | Zf | 1.00E-02 | 0.0407 | 12.59 | 1.23 |

| | | | | | |
|---|---|---|---|---|---|
| ERG | ETS | 1.00E-02 | 0.0419 | 19.28 | 1.17 |
| Foxa3 | Forkhead | 1.00E-02 | 0.0451 | 4.00 | 1.49 |
| ELF1 | ETS | 1.00E-01 | 0.0471 | 4.90 | 1.42 |
| HIF-1b | HLH | 1.00E-01 | 0.0471 | 10.99 | 1.25 |
| EBF | EBF | 1.00E-01 | 0.05 | 3.00 | 1.58 |
